# Metabolic STAMP reveals GPCR signaling networks that program β-cell insulin secretion

**DOI:** 10.1101/2025.10.03.680349

**Authors:** Mohammad Ovais Aziz-Zanjani, Rachel E. Turn, Yan Hang, Anushweta Asthana, Leilani E. LaBrie, Mohammadamin Mobedi, Lucy Artemis Xu, Amanda Ramste, Michael Krawitzky, Joshua W. Knowles, Seung K. Kim, Peter K. Jackson

## Abstract

Pancreatic islet β cells integrate glucose and hormonal cues to control insulin secretion through spatially and temporally organized phosphorylation networks in health and diabetes. Here, using Metabolic STAMP (Synchronized Temporal-Spatial Analysis via Microscopy and Phosphoproteomics), we combine time-resolved phosphoproteomics, imaging, and kinase inhibition in mouse β cells and human islets to map stimulus-specific GPCR signaling pathways. Metabolic STAMP reveals that GLP1-R and FFAR4 engage distinct, compartmentalized kinase programs, including GLP1-R-biased ERK activation, receptor-specific cAMP-PKA domains, and a phospho-ATAT1/HDAC6 node that differentially modulates microtubule acetylation and insulin secretion during GSIS based on stimulation conditions. These GPCR-responsive phospho-signatures and microtubule remodeling patterns are substantially conserved in human islets. Together, our data define an integrated, compartmentalized signaling architecture linking metabolic GPCR inputs, organelle remodeling, and insulin secretion, and provide a β-cell phosphoproteomic resource that connects dynamic signaling nodes to human genetic risk and potential therapeutic targets.

## INTRODUCTION

Pancreatic islet β cells dynamically respond to multiple physiological cues to enhance insulin secretion in response to glucose, including nutrients, neurotransmitters, and hormones. This heightened sensitivity is largely orchestrated by post-translational modifications (PTMs), notably phosphorylations, which act as rapid and reversible regulators of insulin output. The transient nature of phosphorylations, controlled by kinases and phosphatases, tunes the rapid acceleration of insulin secretion in response to glucose within minutes, as well as its timed cessation when glucose levels drop. Previous groups^1–5^ have performed phosphoproteomics of different stimuli in β cells, yet the broad cellular functions of specific protein (de)phosphorylation events remain largely undefined.

Once triggered by glucose, β cell insulin secretion is further modulated by G protein-coupled receptors (GPCRs), which integrate signals from circulating factors. Among these, FFAR4 (Free Fatty Acid Receptors 4) and GLP1-R (Glucagon-Like Peptide-1 Receptor) have been identified as pharmacological targets^6–9^ for regulating Glucose Stimulated Insulin Secretion (GSIS). These receptors sense extracellular cues and transduce them into intracellular signals via post-translational modifications (PTMs) targeting organelle dynamics. Tightly regulated organelle dynamics are crucial for coordinating GSIS. For example, recent research highlights primary cilia as critical signaling hubs that organize GPCR activity in both α and β cells.^10^ Our recent analysis identifies multiple GSIS-regulating ciliary GPCRs that signal through primary cilia, including the ω-3 fatty acid receptor FFAR4^6^ among others. Despite extensive studies on these individual GPCRs to understand mechanism, how multiple GPCRs activate similar or complementary signaling pathways to enhance insulin secretion remains unclear. Especially pivotal to addressing this question is to study the mechanisms by which these signals are regulated in time and space on the order of transient PTMs. Here, we focus on how multiple GPCRs modulate compartmentalization of kinase activity and phosphorylations to different organelles, thus coordinating β-cell responses to metabolic cues. These phosphorylations have the potential to regulate diverse, context-specific features of protein function, including protein localization, binding partner affinity, structural conformation, and enzyme activity.^11^ The majority of phosphorylation sites in islet β cells that have been identified via bulk phosphoproteomics have no known function or linkage to specific signaling pathways.^12^

To address this gap, we developed Metabolic-STAMP (**Fig. S1**) to identify and link GPCR-activated kinases to substrates and to map substrate dynamics during β cell simulation.^13^ Given recent advances in linking ’consensus’ protein sequences of phosphorylation (consensus phosphorylation patterns: CPPs) to specific kinases,^14^ we determined the kinase-CPP pairings that are activated during GSIS, including phosphosites linked to JNK, CDK, PKA, and the CAMK family. We connected phosphorylations to specific “timestamps” and cellular phenotypes using high-resolution microscopy of key cellular markers.^13^ Through these integrative approaches, we *visualized* and quantified critical cellular dynamics of microtubules, cilia, mitochondria, and vesicular trafficking and linked these dynamics to CPP-targeted phosphorylations that regulate β cell function. Metabolic-STAMP revealed differential regulation of microtubule acetylation in GPCR versus glucose stimulation alone in both mouse pancreatic β cells and human islets. This study provides a blueprint of specific GPCR signaling networks and regulatory features that control insulin secretion, and unique phospho-biomarkers to assess β cell function. **As a Resource, this study delivers (i) a β-cell phosphosite atlas (mouse and human), (ii) compartment-resolved UMAPs, (iii) kinase–substrate predictions supported by inhibitor perturbations, and (iv) organelle imaging datasets directly linked to phosphosignatures.** This study provides a roadmap for identifying specific GPCR signaling networks and regulatory features that control insulin secretion and offers a broad panel of phospho-biomarkers to assess β cell signaling.

## RESULTS

### Quantitative phosphoproteomics maps stimuli-specific subcellular signaling dynamics in β cells

We have developed a new protocol called Spatial-Temporal Analysis by Microscopy and Proteomic (STAMP) that correlates detailed microscopy of cellular endophenotypes to phosphoproteomic changes in timed kinetic analysis following a signaling event.^13^ We proposed that Metabolic STAMP could deconvolve time-dependent signaling- and organelle-based mechanisms directing GPCR regulation of insulin secretion.^15, 16^ Initially, we studied the glucose-responsive murine β cell line MIN6 using fragmentation-based phosphoproteomics with a timsTOF-HT mass spectrometer (Methods).^13^ Later, we studied primary human islets (see below). We mapped phosphorylations 10 minutes (min) after cells were exposed to 5 conditions: low glucose (’LG’, 2.8 mM), high glucose (’HG’, 17.8mM), HG + GLP1-R agonist Exendin4 (’HG+Ex4’), HG + FFAR4 agonist TUG891 (’HG+TUG891’), or HG + Ex4 + TUG891 (’Combo’) (see Methods). Previous studies^1^ demonstrated that maximal changes in β cell phospho-signatures occurred ∼10 min after glucose stimulation. In these 5 conditions, we also measured insulin secretion and visualized subcellular organelles by immunofluorescence at 5 and 20 min to assess phenotypes before and after capturing phospho-signatures. As expected,^6, 17^ HG+Ex4 or HG+TUG891 enhanced insulin secretion from MIN6 cells ∼2-fold relative to HG alone, whereas HG+Ex4+TUG891 (Combo) further amplified insulin release within 30 min after stimulation (3-4-fold; **Fig. 1A**). We postulate that dual GPCR agonism leads to coordination of unexplored regulatory mechanisms, stimulating release of pre-formed insulin granules.^18^

**Fig. 1:**
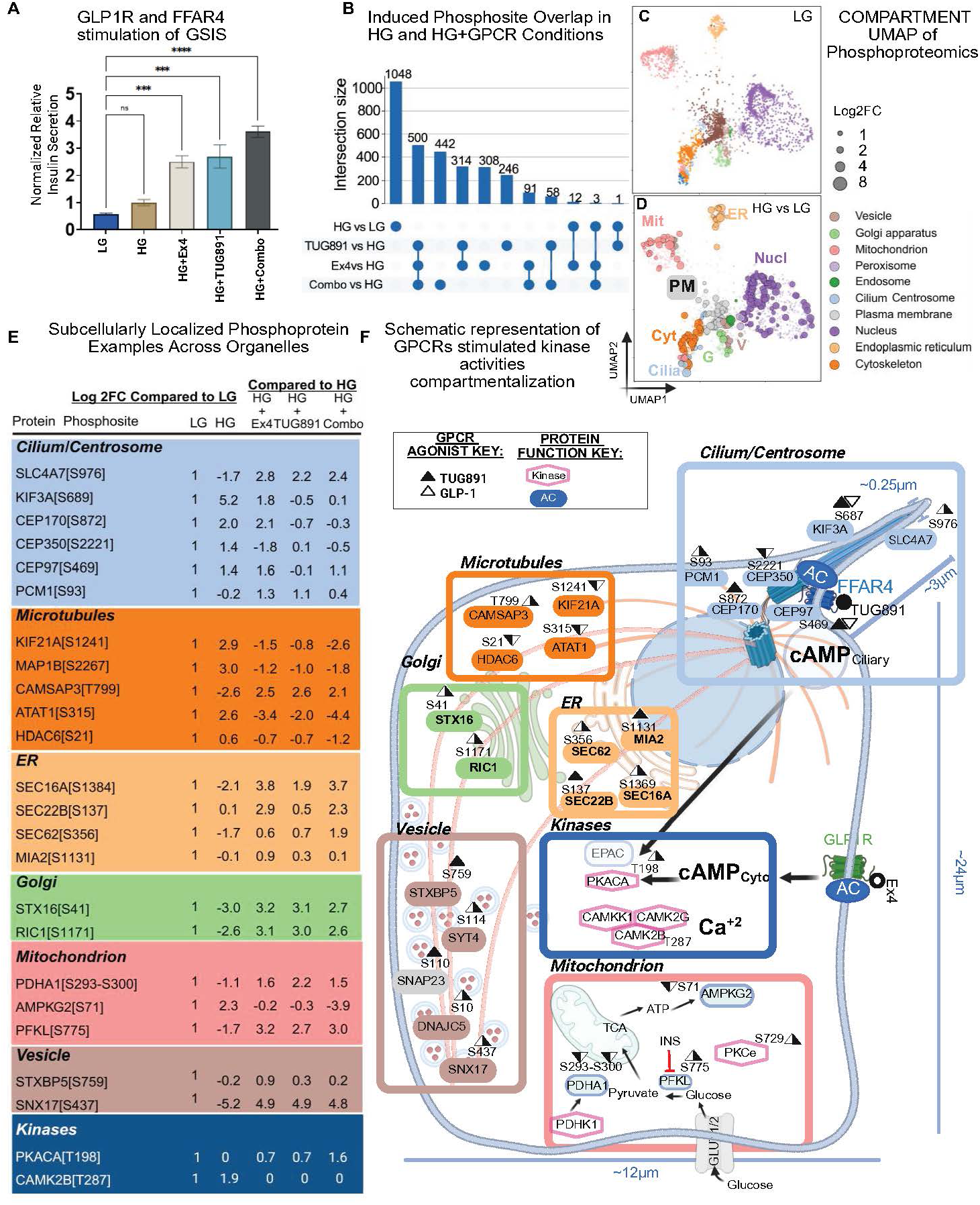
Mapping the differential phosphoproteomics of GSIS with and without co-stimulation with GPCR agonists. (**A**) Normalized relative insulin secretion levels were determined via luciferase assay was measured in mouse pancreatic β cell line MIN6-6, either at low glucose (2.8mM) or treatment with combinations of high glucose (17.8mM), TUG891 (100μM), or Exendin4 (20nM). Cells were serum and glucose starved for 1hr before treatment, and samples were collected at 30 min after treatment. N=3. Error bars= SEM. Statistical significance was assessed via One-Way ANOVA, with \**P* = 0.05; \*\**P* = 0.01; \*\*\*\**P* = 0.0001. Plots were generated using GraphPad Prism software. (**B**) Upset plot of overlapping phosphorylation events induced in different GPCR stimulation conditions or glucose alone in MIN6 cells (>2-fold increase, p<0.001). (**C-D**) UMAP COMPARTMENT analysis reveals differential effects of low versus high glucose on the cellular phosphoproteomic landscape. Circle size indicates log2 fold change, circle color represents the known subcellular compartment where that protein functions. The first panel shows baseline levels of all detected phosphorylation sites, and the subsequent panels display these sites with circle sizes increased for phosphosites that show marked changes in abundance under each stimulation condition. (**E**) Examples of changes in abundance of pathway-specific phosphorylations during GSIS. Differences are reported as log2 fold change compared to low glucose conditions. (**F**) Induced phosphorylation of proteins associated with altered GSIS and important for the function of specific subcellular compartments. These include cilia and centriole (blue), microtubule and kinesins (orange), Golgi and ER (green), secretory machinery (yellow). Proteins previously associated with GPCR regulatory machinery (navy) and glucose metabolism (grey) are noted. Representative phosphorylations are shown here in a β cell model, with key cellular compartments marked. Arrowheads indicate whether it is an increase or a decrease in abundance and which GPCRs/stimulations induce the change (black: GLP1-R; white: FFAR4).

Following 10 min exposure to HG, HG+Ex4, HG+TUG891, and Combo stimulation, compared to LG controls, we detected >17,000 reproducible, high-confidence phosphorylation sites on >3,700 proteins (**Table S1**). The majority of these phosphorylations have not been previously linked to function or localization in β cells. Most phosphosites (14,794 of 17,567) were shared in multiple conditions, although ∼10% were unique to HG, and another ∼9% appeared only with GPCR activation. To pinpoint important regulatory changes, we quantified stimulus-dependent dynamics of these phosphorylations. Phosphorylation changes were defined as proteins exhibiting at least a two-fold increase or decrease with *P*<0.001 on identified phosphorylation sites. After 10 min HG treatment, 1048 phosphorylation events were elevated compared to LG (**Table S1**)- most of these changes were not observed after stimulation with either Ex4 or TUG891 alone. Similarly, 1971 phosphorylations were uniquely induced by either TUG891, Ex4, or Combo, but not by HG alone. Among GPCR-induced phosphorylations, 963 were common across more than one condition, in part reflecting common pathway activation; one prime example of this was cAMP-PKA pathway activation. However, ∼50% of induced phosphorylations were unique to either the TUG891, Ex4, or Combo treatment. Thus, of the phosphorylations revealed during stimulation of insulin secretion, the majority were specifically induced by GPCR-mediated signaling (**Fig. 1B, Table S1**). In addition, >800 sites on 667 proteins that were induced by either HG, HG+Ex4, or HG+TUG891 conditions were instead attenuated after Combo treatment (**Fig. S2A**, **Table S1**).

### Subcellular compartments show differential phosphoprotein regulation during β cell GSIS

To localize GSIS-associated phosphoproteins identified by Metabolic STAMP, we used COMPARTMENTS,^19^ a database for assigning proteins to 10 subcellular compartments, then compared phosphoprotein distributions across treatments (**Fig. 1C-D, Fig. S2B-F; Table S2)**. UMAP analysis resolved ∼3,700 phosphoproteins into these 10 compartments, revealing differential effects of HG versus HG+GPCR activation on β-cell endophenotypes during GSIS. These data suggest that GPCR-driven signaling engages compartment-specific phosphorylations, revealing spatial selectivity of kinases activated downstream of β-cell GPCRs **(Fig. 1E-F, Fig. S2B-F)**.

Quantitative phosphoproteomics identified GPCR agonist-induced phosphorylations of multiple substrates, including kinases, phosphatases, microtubule-associated proteins, ciliary proteins, kinesins, mitochondrial proteins, secretory pathway proteins, and protein transport factors, most not previously reported in β cells (**Fig. 1E-F**). We observed changes in the abundance of phosphosites, with induction or suppression of specific phosphosites depending on treatment conditions (**Fig. 1E**). For example, phosphorylations of known, crucial regulators of cilia and centrosome, including kinesin KIF3A on [S687] and centrosomal protein CEP97 on [S469], were differentially regulated by TUG891 versus Ex4 treatment (**blue panel, Fig. 1E**). In both cases, TUG891 stimulation suppressed phosphorylation to levels seen in LG, while Ex4 increased CEP97[S469] phosphorylation. KIF3A[S687] fits a PKA consensus phosphorylation site and is known to promote vesicular cargo loading and motor function in neurons,^20–22^ but the role of this site in β cells has not been elucidated. Phosphorylation of the microtubule cytoskeletal machinery showed strong *opposite* fluctuations in abundance upon HG+GPCR stimulation compared to HG stimulation alone (**orange panel, Fig.1E**). These included GPCR agonist-dependent de-phosphorylation of MAP1B[S2267] and microtubule growth regulator kinesin KIF21A[S1241], which are in high abundance upon HG stimulation but suppressed by Ex4, TUG891, and Combo treatments. Conversely, phosphorylation of Calmodulin-regulated spectrin-associated protein 3 CAMSAP3[T799] is strongly suppressed by HG alone but is increased at least two-fold by stimulation of GLP1-R or FFAR4 (**orange panel, Fig. 1E**). CAMSAP3 is a critical regulator of minus-end, non-centrosomal microtubules that prevents depolymerization and stabilizes microtubules.^23^ For other targets, like SEC22B[S137] (a SNARE protein family member that regulates ER-to-Golgi vesicle trafficking), we observed modest increases of phosphorylation after HG, which was further increased with exposure to Ex4, TUG891, and Combo treatments (**tangerine panel, Fig. 1E**). While regulation of this phosphosite has not been previously noted in β cells, prior studies of hepatocytes reported that SEC22B[S137] phosphorylation is induced after stimulation with glucagon to regulate binding partner selectivity.^24^ Together, these quantitative phosphoproteomics provide a new roadmap to decode complex signaling networks underlying insulin secretion. These data further reflect mechanisms involving signaling synergy, negative feedback, and pathway specificity between different glucose and GPCR stimulation conditions, indicating how these different pathways can coordinately tune insulin secretion.

### Phosphosite mapping unveils stimuli-specific activation of compartmentalized PKA and ERK signaling in β cells

Kinases like Protein Kinase A (PKA) and Extracellular Signal-Regulated Kinase (ERK) are known regulators of β cell GSIS, as shown by prior loss-of-function and pharmacologic studies,^25, 26^ but how these kinases respond to distinct stimuli to implement signaling specificity and flexibility in β cells remains incompletely understood. To fill this knowledge gap, we extracted phosphorylation sites from our phosphoproteomics dataset and overlaid them onto a public signaling network (PhosphoPlus®^27^), tracing differential phosphorylations triggered by HG versus HG+GPCR stimulation. We then used a kinase inhibitor screen (see below) to test these correlations. This focused approach revealed phosphorylation of known and unrecognized sites phosphorylated by PKA, ERK, and other pathways (**Fig. 2**), thus unveiling regulation of multiple signaling network roles in GSIS.

**Fig. 2:**
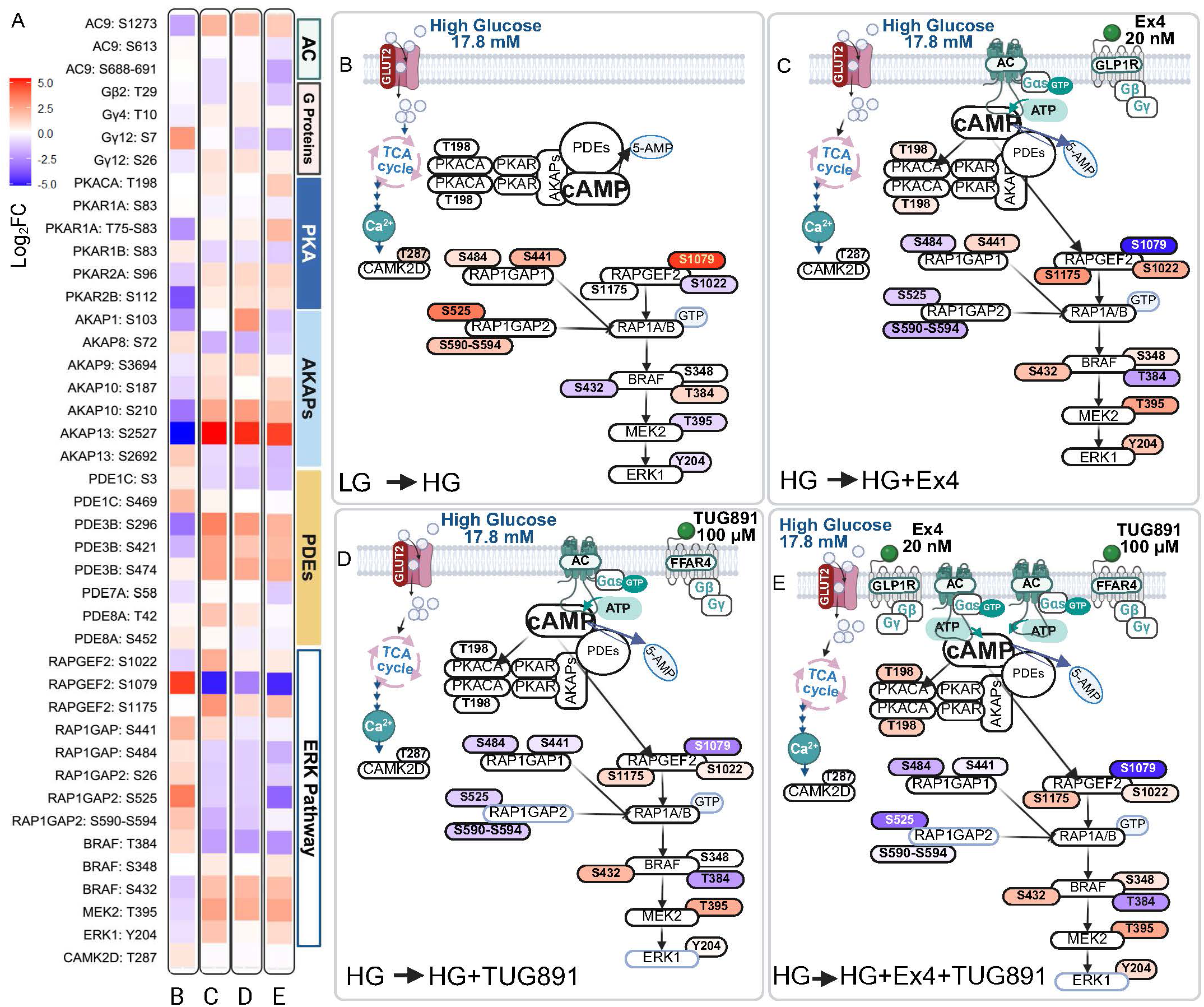
Map of the ERK pathway phosphoproteomic signatures of MIN6 cells in different GSIS regulatory conditions. Here, we highlight key phosphorylations in known signaling pathways and how they are differentially regulated in the context of GSIS with and without GPCR stimulation. **(A)** Heat map showing the log_2_-fold change in phosphorylation abundance of PKA- and ERK-pathway signaling regulators upon stimulation with high glucose and the indicated GPCR agonists for 10 min. Phosphosites are grouped by protein family, as indicated on the right. Phosphorylation levels are reflected according to heat map, with increased abundance in red and decreased abundance in blue. (**B**-**E**) Schematic of ERK phospho–signaling, highlighting which branches of the pathway are engaged under each treatment condition, including (**B**) HG, (**C**) HG+Ex4, (**D**) HG+TUG891, and (**E**) HG+Combo. The ERK signaling phosphoproteomics data reveal how the corresponding phospho–signatures differ. Phosphosites are depicted using the same heat–map color scale as in (**A**), reflecting their relative abundances based on experimental condition.

PKA catalytic and regulatory subunits heteromerize with cAMP-activated signaling complexes, scaffolding protein A-kinase anchoring proteins (AKAPs), and phosphodiesterases (PDEs), which catabolize and inactivate cAMP (formed by adenylate cyclases, AC; **Fig. 2, Fig. S2G**). Together, these signaling complexes and elements orchestrate temporal-spatial regulation of responses in β cells and other cell types.^28, 29^ We detected high levels of phosphorylation of the PKA catalytic subunit at [T198] at high abundance with HG or HG+GPCR stimulation, validating the robustness of our dataset. Additionally, we observed differential isoform-specific phosphorylations of other complex components. In HG, we observed elevated phosphorylation of cytoplasmic Type 1 regulatory subunit PKAR1B[S83], whereas phosphorylation of organelle-anchored Type 2 subunits (*e.g*., PKAR2A[S96] and PKAR2B[S112]) was enhanced by GPCR activation (**Fig. 2A, Fig. S2H-K**). PKAs are known to be locally regulated by distinct AKAP family members in discrete compartments, and we observe notable changes in induced phosphorylation of AKAPs that could activate their precise temporal and spatial signaling regulation (**Fig. S2H-K**). For example, Ex4 and TUG891 augmented phosphorylation of AKAP1, AKAP9, and AKAP10, previously reported to localize to mitochondria, Golgi, cytoskeleton, and plasma membrane, respectively ^30, 31^ (**Fig. S2H-K)**. In contrast, HG selectively induced phosphorylation of nuclear AKAPs (**Fig. S2H-K**). We also noted localized phosphorylations of phosphodiesterases (PDEs), which are known to ’tune’ PKA activity by locally dampening cAMP gradients.^32^ HG induced phosphorylation of cilium-associated PDE1C, whereas GLP1-R and FFAR4 stimulation led to phosphorylation of membrane-bound PDE3B^33, 34^ at multiple phospho-sites (**Fig. S2H-K**). By contrast, FFAR4 activation uniquely elevated mitochondrial AKAP1[S103] while reducing phosphorylation of mitochondrial PDE8A[S452] (**Fig. S2H-K**), a unique PKA activation pattern not observed with either glucose or GLP1-R stimulation. Given the link between rapid FFAR4-dependent ciliary cAMP activation and ciliary elongation (see below), identifying the key cAMP effectors (like specific AKAPs and PDEs) nominates strong candidate mediators of these events.

In parallel, we observed stimulus-specific activation of the ERK pathway. HG+Ex4, but not HG+TUG891 or HG alone, induced a cascade of stimulatory phosphorylations on GAPGEF2, RAP1, BRAF, MEK2, and ERK[Y204] (**Fig. 2B-E**). Immunoblotting (**Fig. S2M**) confirmed that GLP1-R stimulation enhanced phospho-ERK activity compared to FFAR4. Collectively, these results reveal that different stimulus combinations engage distinct components of pathways regulating β cell secretion.

### Kinase-inhibitor screen corroborates phosphoproteomic predictions and reveals regulators of β cell secretion

Our phosphoproteomics nominated multiple kinases as regulators of glucose-stimulated and GPCR-potentiated insulin secretion in β cells. To test functional dependencies, we screened the Gray Lab kinase library of 240 validated kinase inhibitors^35–37^ for effects on insulin secretion (Methods, **Fig. S3**). MIN6 cells were pretreated for 1hr in LG with each inhibitor, then stimulated with Combo treatment (HG+Ex4+TUG891) in the continued presence of inhibitor, followed by measuring insulin secretion after 30 min. Multiple kinase inhibitors suppressed GSIS, including 17 kinase inhibitors or regulators that suppressed Combo-stimulated insulin secretion (**Fig. S3**, **Fig. 3A, Table S3**) by 4-fold or more. These included inhibitors of targeted kinases known to regulate insulin secretion, like PKA, and those not previously linked to insulin secretion regulation, like HEC1.

**Fig. 3:**
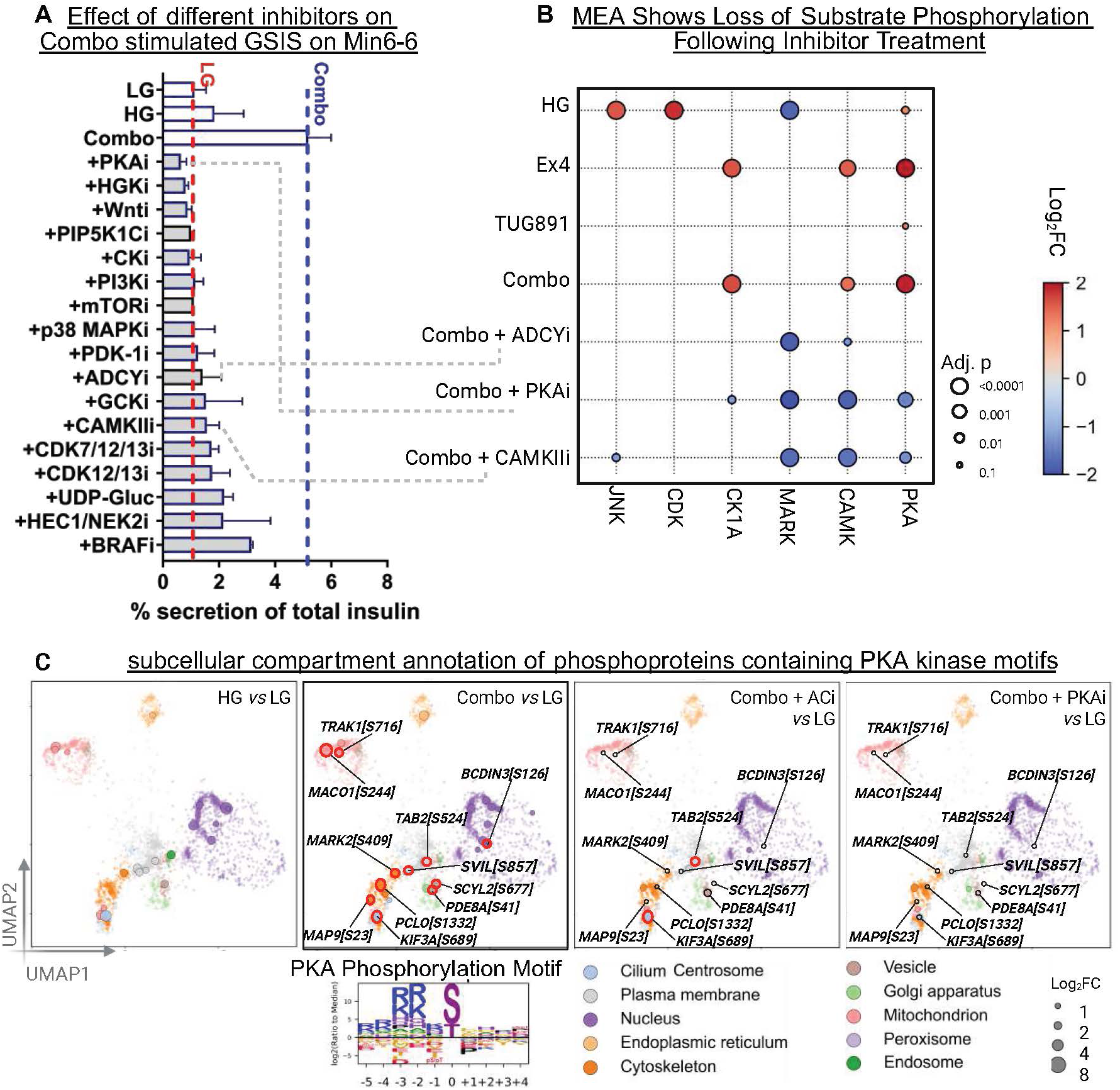
Mapping the context-dependent regulation of kinases during MIN6 GSIS with and without GPCR activation. **(A)** Plot of the effects of different protein kinase and pathway inhibitors (15 kinase inhibitors, ADCY inhibitor, P2YR14 receptor agonist) in combination with HG and Combo treatment on % insulin secretion. Error bar= SD. Note the red dotted line that indicates insulin secretion levels at low glucose and the blue dotted line indicating the baseline insulin secretion with Combo GPCR agonist treatment. **(B)** Dot plot showing the fold change in inferred kinase activity calculated by Motif Enrichment Analysis (MEA). Increasing red coloration indicates increased activity, while increasing blue coloration indicates decreased activity. Dot size reflects the p–value. Each column corresponds to a kinase whose activity was inferred by MEA using PhosphoSitePlus® (see Fig. S2 for workflow), and each row represents a specific treatment condition. Arrows on the left link the conditions in panel (**A**) to their corresponding columns in the dot plot on the right. **(C)** UMAPs showing the subcellular localization of known PKA substrates and the differential impact of stimulatory versus inhibitory treatments on their phosphorylation abundance. On the far right, the legend indicates the compartment associated with each circle color, and circle size reflects the log2 fold change in phosphorylation. Example putative PKA substrates are highlighted that show increased phosphorylation under Combo treatment (second panel) and reduced phosphorylation upon adenylyl cyclase inhibition (third panel) or PKA inhibition (fourth panel). Experimental phosphoproteomics data from Combo samples treated with adenylyl cyclase or PKA inhibitors confirm the inferred putative PKA substrates, as their phosphorylation is selectively reduced under these inhibitory conditions.

To identify kinase dependency of specific β cell phosphosites, we performed phosphoproteomics on MIN6 cells treated with HG+Ex4+TUG891 (Combo), together with inhibitors targeting PKA (H-89), Casein Kinase (Silmitasertib), ADCY (NKY-80), or CAMK (NK-93). Motif enrichment analysis (MEA;^14^ see **Methods)** confirmed loss of putative substrate phosphorylation with inhibitors that also suppressed insulin secretion (**Fig. S3**, **Fig. 3A-B**). For instance, phosphosites enriched in PKA motifs were selectively and strongly reduced after treatment with PKA selective inhibitor H-89, while sites matching CAMKII motifs were strongly reduced after KN-93 treatment (**Fig. 3B, Table S1**). While changes in kinase-selective phosphorylations were expected, our screen also identified target phosphosites for PKA not previously catalogued (*e.g.*, SEC16A[S1384] and RAB3GAP1[S537]) and for CAMKII (*e.g.*, VAMP2[S75], SEC22B[S137]). In summary, PKA and CAMKII substrates calculated using Cantley metrics were further validated by decreased phosphosignals following selective kinase inhibitor treatment in phosphoproteomic analyses, providing strong functional evidence of kinase activity and associated downstream targets. Sites that did not show reduced signal may be attributed to phosphorylation by other kinases that recognize similar motifs.

Unexpectedly, we also observed a subset of PKA and CAMKII-selective phosphorylations suggesting kinase crosstalk. For example, PKA inhibition was sufficient to suppress some CAMKII motif phosphorylations, and vice versa (**Fig. 3B**). Specific substrates, like the microtubule-associated kinase^38, 39^ MARK2[S409], were sensitive to both PKA and CAMKII inhibitors. In parallel, inhibition with ADCY inhibitor (NKY-80) revealed overlap with PKA-dependent sites as well as distinctive targets (**Fig. 3B, C; Table S1**). MEA revealed distinct kinase activities during GSIS beyond PKA and CAMK. Putative MARK kinase substrates were reduced upon glucose stimulation, an effect that was reversed by GPCR activation, with neither PKA nor CAMK inhibition having a significant impact. In contrast, CDK and JNK substrate phosphorylation increased in response to glucose and was attenuated by stimulation of either GLP1-R or FFAR4 (**Fig. 3B**). Notably, a subset of putative CDK substrates included proteins associated with microtubule plus-end networks implicated in insulin secretion hotspots, which exhibited similar regulation patterns^40^ (**Fig. S2L**). Together, these data show that GPCR activation differentially regulates MARK, CDK, and JNK substrate phosphorylation during GSIS, defining stimulus-specific changes in kinase-associated phosphorylation signatures.

We next performed compartment analysis to ask if there was subcellular localization of kinases and ADCY mechanisms regulating GSIS. Our findings revealed that compartments including the microtubule cytoskeleton, cilium, mitochondria, vesicle, and plasma membrane were enriched for candidate PKA-phosphorylated proteins (**Fig. 3C**). Many similar phospho-sites were suppressed by both ADCY and PKA inhibitors, including cytoskeletal components MAP9[S23] and MARK2[S409] and plasma membrane protein SVIL[S857]). However, additionally, some phospho-sites appeared attenuated only by PKA inhibitor but not ADCY inhibitor, including ciliary KIF3A[S689] and plasma membrane TAB2[S524]. These kinase inhibitor studies support our β cell phosphoproteomics results and demonstrate both specificity and convergence of signaling pathways downstream of HG versus GPCR stimulation.

### Stimulus-specific remodeling of β cell organelles

To test whether phosphorylation dynamics translate into intracellular remodeling that could direct β cell function, we performed high-resolution confocal microscopy of key compartments to reveal context- and time-specific fluctuations in phospho-signaling in cilia, microtubules, mitochondria, and Golgi apparatus. We observed examples of both pathway-specificity and synergy in each compartment. For example, we detected dynamic, stimulus-specific organelle remodeling. HG and HG+GPCR stimulation led to reduced residence of β-COP (a marker for COP-I vesicles) in the *cis*-Golgi, marked by GM130, but corresponding increases in vesicular staining (**Fig. 4A**). This was consistent with an increase in remodeling of the Golgi and mobilization of secretory machinery in response to increased metabolic demand upon glucose treatment. While β-COP staining after HG alone or HG+TUG891 treatment were similar, HG+Ex4 stimulation led to an increase in β-COP cytoplasmic staining within 5 min (**Fig. 4A**). Establishment of transient mitochondrial-Golgi contacts was markedly delayed after HG+Ex4 stimulation, compared to HG or HG+TUG891 (**Fig. 4B**).

**Fig. 4:**
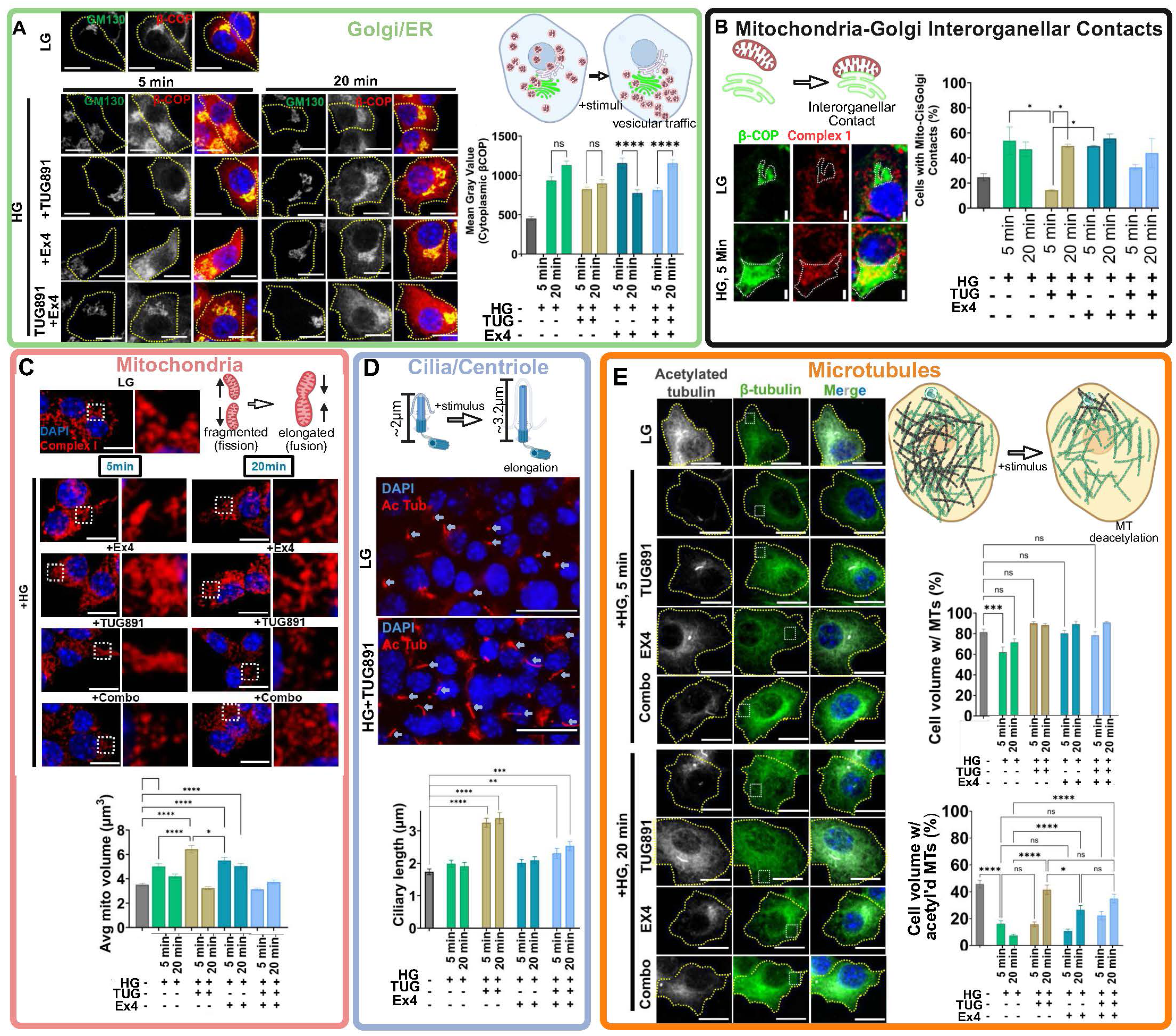
Differential GPCR and glucose stimulations lead to differential effects on cellular compartments based on fixed cell imaging. Changes in key cellular compartments with different stimulation conditions were assessed via immunofluorescence and confocal microscopy at 60x magnification. Samples were collected at 5min and 20min after stimulation. The following compartments were assessed, with representative images and image quantification shown for each: (**A**) Golgi and vesicular traffic machinery morphology and spreading with dotted lines indicated cell borders (magenta: β-COP; cyan: GM130; blue: DAPI) (scale = 10µm), and quantification of mean gray value cytoplasmic β-COP was assessed for each condition (far right); (**B**) mitochondria-Golgi inter-organellar contacts, with dotted lines outlining the Golgi border to reflect whether mitochondrial are excluded (left, low glucose) or overlapping with Golgi (right, high glucose 5min) (blue: DAPI; green: β-COP; red: Complex-I) (scale = 10μm); quantification on left showing what percentage of cells in each condition show evidence of overlap with Golgi, with error bar= SEM; (**C**) mitochondrial morphology, with blown up insets showing representative regions highlighting the extent to which mitochondria are tubular/fragmented for each condition (green: Complex 1) (scale = 10µm), with average mitochondrial volume being assessed for each condition (below); (**D**) ciliary lengthening (red: acetylated tubulin; blue: DAPI) (scale = 20µm), with graph quantifying average length of cilia in each condition; (**E**) total microtubule density (green: β-tubulin) versus acetylated microtubules and yellow lines marking cell borders (white: acetylated tubulin; blue: DAPI) (scale = 10µm); differences in density were measured as % cell volume encompassed by total tubulin (upper right) versus acetylated tubulin (lower right); Experiments were performed in biological replicate, with ∼100 cells per replicate except for mitochondrial volume which was >2000 mitochondria per condition and ∼20 cells per condition for β-COP. Error bars= SEM. Statistical significance was assessed via One-Way ANOVA, with \**P* = 0.05; \*\**P* = 0.01; \*\*\*\**P* = 0.0001. Plots were generated using GraphPad Prism software.

Across treatment conditions, we also examined morphology of mitochondria and cilia, highly dynamic organelles whose morphology changes in response to cellular nutrient status.^41, 42^ Complex I is a mitochondrial outer membrane marker (**Fig. 4C**) and key regulator of the electron transport chain. We observed an increase in average mitochondrial volume (elongation) at 5 min after HG or HG+GPCR stimulation, consistent with increased mitochondrial fusion or reduced mitochondrial fission.^43^ This increased volume resolved by 20 min after HG or HG+TUG891 stimulation; by contrast, mitochondrial elongation after HG+Ex4 was prolonged, while HG+Combo treatment prevented increased mitochondrial volume (**Fig. 4C**). This spectrum of responses appeared to correlate with selective phosphorylations of mitochondrial elements, including increased phosphorylation of the PKA substrate Trafficking kinesin-binding protein TRAK1[S716], a critical regulator of mitochondrial motility (**Fig. 3C**). For primary cilia, HG+TUG891 stimulation strikingly triggered rapid and pronounced ciliary elongation within 5 minutes, which persisted for over 20 min (**Fig. 4D**), whereas no significant changes were observed in HG or HG+Ex4 groups. Cells treated with the Combo condition demonstrated reduced ciliary lengthening compared to HG+TUG891, suggesting competition or negative cross-regulation of GLP1-R on FFAR4 signaling (**Fig. 4D**). These findings align with our phosphoproteomic data indicating enrichment of FFAR4-regulated ciliary phosphoproteins, including the sodium-bicarbonate co-transporter^44^ SLC4A7[S976] (**Fig.1D**). Thus, HG and distinct GPCR signals stimulated specific intracellular machinery and induced organelle remodeling during potentiation of insulin secretion, underscoring stimulus-specific compartmentalization of signaling during β cell GSIS.

### Phosphorylation of ATAT1 and HDAC6 and regulation of β cell microtubule remodeling

We noted a distinct, coherent pattern of β cell microtubule remodeling and PTMs during GSIS. The abundance of total assembled tubulin (assessed by immunostaining of β tubulin) only showed modest fluctuation, while HG led to significant microtubule disassembly, especially along the cell periphery (**Fig. 4E**). This was consistent with previous reports suggesting that peripheral microtubule disassembly after HG is important for insulin secretion.^45^ Thus we hypothesized that microtubule disassembly might be linked to destabilization by deacetylation, as in primary cilia.^46–48^ Staining for acetylated tubulin revealed striking differences between Ex4 and TUG891 stimulation (**Fig. 4E**). HG stimulation led to persistent microtubule deacetylation after 5 and 20 min. On the other hand, HG+TUG891 and HG+Ex4 treatment led to stimulation of a conserved pool of acetylated microtubules localized to the centrosome and Golgi, respectively. After HG+Ex4, the prolonged acetylated microtubule pool appeared localized to *cis*-Golgi, consistent with previous work demonstrating that the Golgi microtubule organizing center (MTOC), not the centrosome, serves as the primary microtubule nucleation site in β cells, in contrast to undifferentiated and/or proliferating cells where the centrosome is the primary MTOC.^45, 49, 50^ While HG+TUG891 stimulation also decreased acetylated microtubules by 5 min, we observed rapid recovery by 20 min. HG+Combo treatment led to reduced microtubule deacetylation at 5 min compared to the other treatments and recovery of acetylated microtubules by 20 min. Thus, we predicted that β cell microtubule acetylation modulates phosphorylation of key regulatory enzymes.

ATAT1 and HDAC6 are enzymes that regulate microtubule stability by controlling the reversible acetylation of α-tubulin on Lysine 40 (**Fig. 5A**).^51^ Our phosphoproteomics analysis identified ATAT1[S315] and HDAC6[S21] as prominent β cell cytoskeletal phosphosites (**Fig. 5C-F**), with phosphorylation increased most by HG alone compared to HG+GPCR signaling (**Fig. 5D, F**). To date, the function of these phosphosites in regulating microtubule acetylation is unknown.^52, 53^ During stimulation of β cells by HG+Ex4+TUG891, exposure to the highly selective HDAC6 inhibitor Ricolinostat significantly blunted insulin secretion (**Fig. 5G; Fig. S4A**), supporting HDAC6 as a previously unrecognized regulator of β cell secretion.

**Fig. 5:**
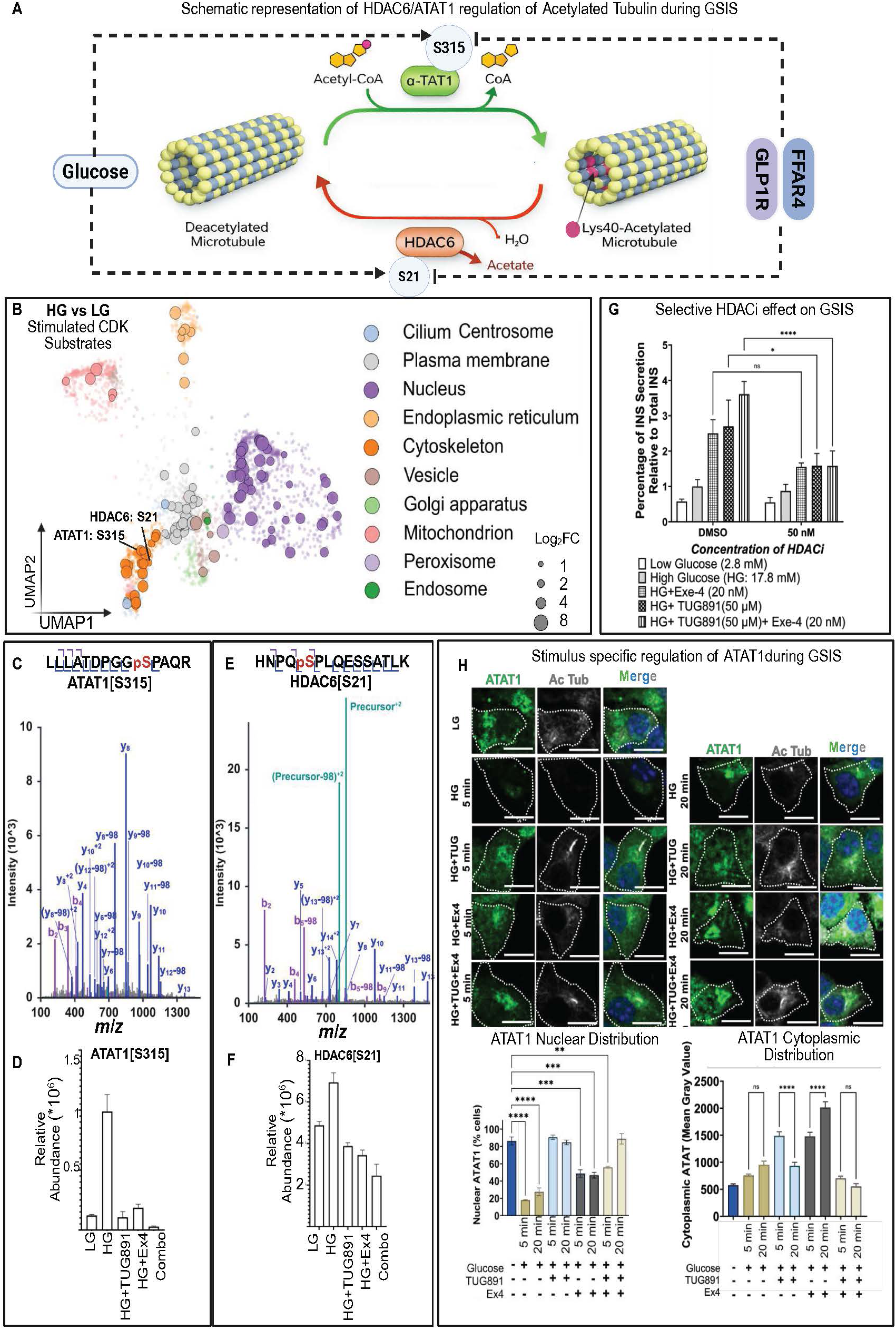
Differential HDAC6 and ATAT1 phosphorylation serve a pivotal role in orchestrating context-specific microtubule stability and dynamics during GSIS. (**A**) Schematic illustrating the roles of ATAT1 and HDAC6 in regulating microtubule acetylation. ATAT1 catalyzes acetylation of α–tubulin at [K40], whereas HDAC6 deacetylates α–tubulin. Two key phosphorylation sites, ATAT1[S315] and HDAC6[S21], are highlighted as putative regulators of this cycle, with high glucose increasing these phosphorylation events and GPCR co–stimulation suppressing them. Figure created with BioRender. (**B**) UMAP-based compartment analysis of putative CDK phosphorylation sites shows increased phosphorylation under HG (10 min) compared to LG in MIN6 cells. Within the high-glucose CDK substrate set, we highlight phosphosites on the key acetylation/deacetylation regulators HDAC6[S21] and ATAT1[S315], both of which are conserved in humans. (**C, E**) Representative spectra of ATAT1[S315] (**C**) and HDAC6[S21] (**E**), and phosphopeptides as generated in Skyline. Above each spectrum, the peptide sequence is shown with the phosphorylated residue highlighted in red and annotated with a preceding “p.” The observed b- and y–ion series support localization of the phosphate to the indicated residue and rule out phosphorylation of adjacent sites. (**D, F**) Plots showing the relative abundance (x10×6) of ATAT1[S315] (**D**) and HDAC6[S21] (**F**) in the following different GSIS treatment conditions in MIN6 cells, 15 minutes after stimulation: LG, HG, HG+TUG891, HG+Ex4, and HG+Combo. Error bars= SD, N=3. (**G**) Plot measuring the effects of a selective HDAC6 inhibitor treatment (5nM Ricolinostat (ACY-1215)) on the percentage of insulin secretion, as measured by luciferase assay in MIN6-6 cells. Cells were grown in low glucose and preincubated either with DMSO or HDAC6i. Samples were treated with one of the following conditions for 30 minutes before harvesting supernatant (without washout of DMSO or HDAC6i): LG, HG, HG+TUG891, HG+Ex4, or HG+Combo. Error bars=SD, N=3. (**H**) Analysis of ATAT1 localization via immunofluorescence of MIN6 cells at different time points after stimulation of GSIS conditions. Above are shown confocal z-stack images (60x magnification) were collected, with green= ATAT1, white= acetylated tubulin, blue= DAPI. Scale= 10μm. Below are plots showing quantification of ATAT1 localization at nucleus (left) versus cytoplasm (right) under different treatment conditions. Error bars= SEM. Statistical significance was assessed via One-Way ANOVA, with \**P* = 0.05; \*\**P* = 0.01; ***P=0.001; \*\*\*\**P* = 0.0001. Plots were generated using GraphPad Prism software.

Metabolic STAMP revealed distinct subcellular dynamics of ATAT1 following specific GSIS stimulatory conditions. Prior studies have linked ATAT1[S315] phosphorylation to nuclear exit.^44, 52^ In baseline LG, ATAT1 was broadly distributed in MIN6 cell nuclei and cytoplasm (**Fig. 5H**), as well as the *cis*-Golgi apparatus (marked by GM130: **Fig. S4B**). HG triggered nuclear export of ATAT1 within 5 min, persisting through 20 min (**Fig. 5H**, bottom panel, **Fig. S5**). HG+Ex4 stimulated modest nuclear export, while HG+TUG891, and HG+Combo, elicited little change (**Fig. 5H**, bottom panel, **Fig. S5**). GPCR stimulation also led to differential ATAT1 distribution outside the nucleus, with Ex4 leading to increased ATAT1 Golgi localization, while TUG891 led to durable ATAT1 cytoplasmic distribution (**Fig. 5H, Fig. S4C**). Collectively, these results demonstrate that glucose and GPCR agonists engage ATAT1 and HDAC6 through distinct phosphorylation events to orchestrate β cell microtubule remodeling, thereby linking cytoskeletal dynamics and insulin secretion.

### Microtubule acetylation dynamics during GSIS recapitulate in human islet β cells

To address whether patterns of regulated microtubule acetylation and assembly were also observed in pancreatic islets, we performed Metabolic STAMP on human cadaveric donor islets (see **Table S4**) exposed for 5, 15, and 20 min to LG, HG, or HG+Ex4+TUG89. Like in MIN6 cell studies, we observed HG triggered deacetylation of microtubules in Insulin^+^ human β cells within 15 min (**Fig. 6A-B**), while stimulation in HG with Ex4 and TUG89 clearly attenuated microtubule deacetylation (**Fig. 6C**). In contrast, Glucagon^+^ human α cells appeared to be largely devoid of acetylated microtubules in all conditions tested (**Fig. 6A-C**, **Fig. S6** green). Together, these data revealed β cells in intact human islets exhibit stimulus-specific microtubule dynamics similar to those in the mouse β cell line (**Fig. 6D**).

**Fig. 6:**
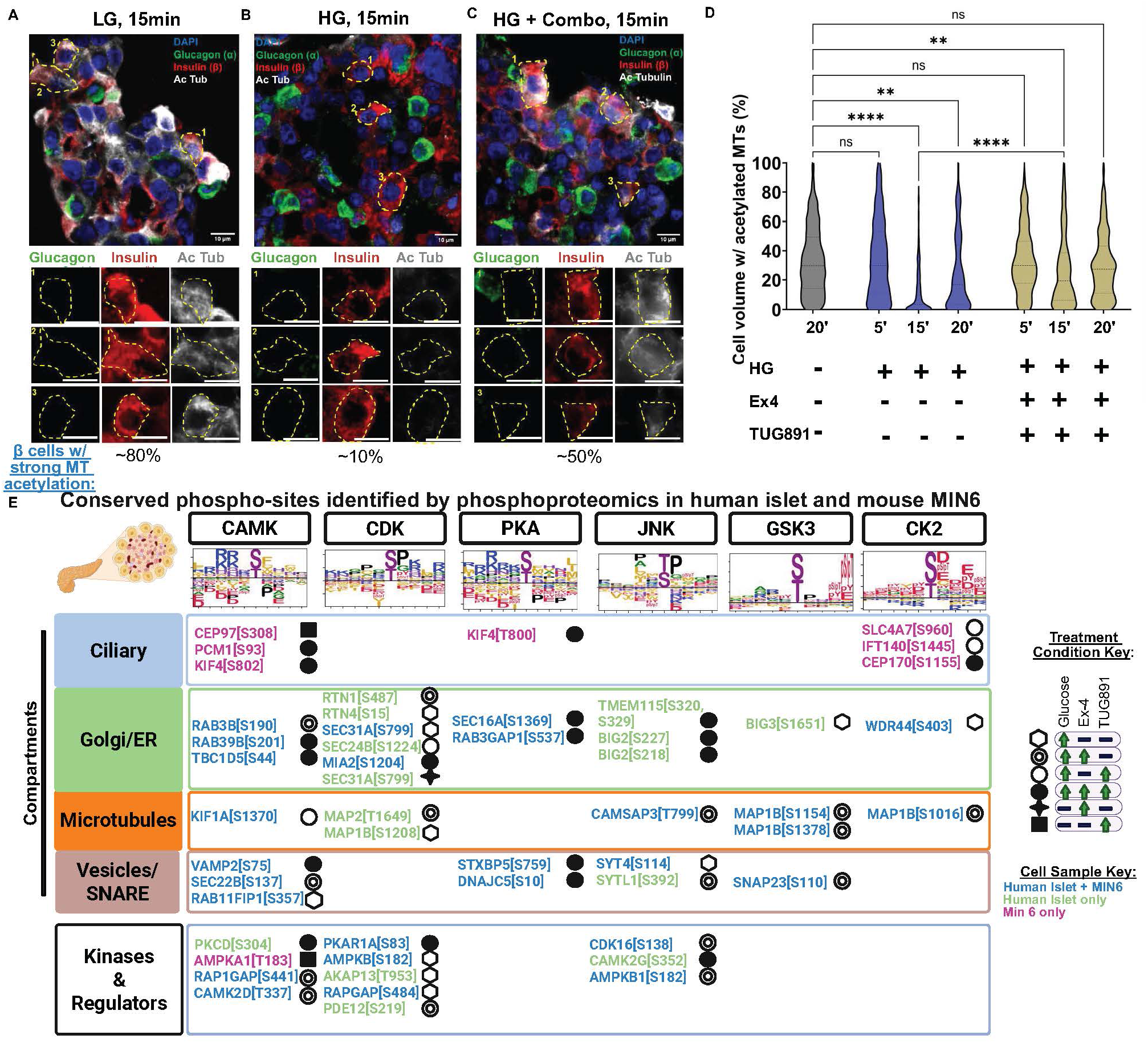
Acetylated microtubule dynamics during different stimulations of GSIS are recapitulated in human cadaveric islets. (**A-C**) Confocal microscopy (60x magnification, oil, z-stacks with 0.28µm per slice) was performed of each experimental condition. Representative images are shown here of the 15min time point for LG vs HG vs HG + Combo (TUG891 + Ex4). DAPI= blue, glucagon= green, insulin= red, acetylated tubulin= white. Scale bar = 10μm for 60x image, scale bar = 5μm for inset. Images were processed using FIJI imaging software. **(D)** Quantitation of the average percent cell volume that contains acetylated microtubules was performed across different experimental conditions and time points, with islets from 3 patient samples (male, between ages 18-50). See Methods/Table S4 for details about human patient samples. Individual dots represent data percentages for individual β-cells (as detected by insulin staining), and the middle bar reflects the mean for that test group. Data were processed in GraphPad Prism, with One-Way ANOVA as statistical analysis; \**P* = 0.05; \*\**P* = 0.01; \*\*\**P*=0.001; \*\*\*\**P* = 0.0001. **(E)** Human cadaveric islets and mouse MIN6 cells show similar changes in compartmental phosphoproteomic signatures following different GSIS treatment conditions. Table showing representative phosphorylations detected from human pancreatic islet (3,000 IEQ per condition). Font color indicates which GSIS phosphorylations are shared between human cadaveric islets and mouse MIN6 stable cell lines (pink: MIN6 only, black: human islet only, blue: shared). Phosphorylations are sorted by row based on known cellular compartment and sorted by column based on predicted regulatory kinase (along with the kinase motif analysis below). Dots next to the phosphorylation reference the code on the right, which highlights the pattern of differential phosphorylation abundance for each experimental condition.

### Shared and divergent phospho-signaling programs in human and mouse β cells

To determine if GSIS phosphoproteome dynamics, we found in mouse β cells were conserved in human islets we mapped HG GPCR-induced phosphorylation in human islets donated from previously healthy subjects (**Table S4**). Perifusion assays of Insulin secretion provided evidence of GSIS stimulated by GPCR agonists in these islets (**Fig. S2N**). Based on imaging analysis detailed above, we profiled the phosphoproteome of human islets after 15 min exposure to four conditions: LG, HG, HG+Ex4, and HG+TUG891. We detected >5,000 phosphorylation events on 1,800 proteins (threshold, FDR<0.01), a yield likely reflecting the smaller sampling of human islet cells compared to MIN6 cells. Compared to LG, we observed increased phosphorylation levels in HG, HG+Ex4, and HG+TUG891 (**Table S5, Fig. 6E**) of multiple proteins, including kinases, calcium regulatory factors, Golgi/ER proteins, MT-associated proteins, and secretory vesicle factors/SNAREs. Remarkably, the majority of phospho-sites induced by HG or GPCR agonists were substantially conserved in the phosphoproteome from similarly-stimulated MIN6 cells (**Fig. 6E**: blue font), including factors encoded at loci containing causal genetic variants linked with diabetes or related metabolic traits by GWAS (like *SEC16A*).^54, 55^ Phosphorylation at some phospho-sites like DNAJC5[S10] and STXBP5[759] increased in HG and was further increased by the addition of either GPCR agonist (**Fig. 6E**). In other cases, we observed agonist-specific phosphorylation. For example, levels of MAP2[T1649] phosphorylation were induced by HG and in HG+Ex4, but not HG+TUG891. Conversely, elevation of several phosphopeptides was detected in human islets but not MIN6 cells (**Fig. 6E**, green font; **Table S1, S5**). This latter finding may reflect species-specific substrates, differing β cell responses in whole islets versus cultured β cell lines, phosphoprotein dynamics reflecting non-β cells including α cells, or technical dropouts from limiting human islet sample size. Collectively, our phosphoproteomics studies in human islets identified multiple conserved glucose- and GPCR-dependent responses in cells, providing a framework for further analysis of primary human islet cells with Metabolic STAMP.

## DISCUSSION

Here, we used Metabolic STAMP^13, 56^ to integrate powerful strategies to dissect conserved glucose and GPCR signaling mechanisms coordinating GSIS in pancreatic β cells. This revealed thousands of phosphorylations occurring within minutes of β cell stimulation, with discrete signatures based upon stimulus. In parallel, Metabolic STAMP generates a β-cell phosphoproteomic resource that maps kinase consensus motifs, subcellular localization, and stimulus dependence for each site, providing a predictive atlas for future β-cell signaling studies. We further show that these predictions have functional traction. These include a selective pharmacologic inhibitor screen for identifying kinases required for GSIS, inhibitor-phosphoproteomics that were consistent with many predicted PKA, CAMKII, and ADCY substrates, and imaging confirming stimulus-specific remodeling of cilia, mitochondria, microtubules, and Golgi. In this study, our focus was on the effects of glucose alone versus differential GPCR activation (in this case, FFAR4 versus GLP1-R). Our findings revealed both synergistic and attenuated effects of these different stimuli combinations on phosphorylation events during GSIS. A selective pharmacologic inhibitor screen identified kinases important for phosphorylation cascades in β cells, confirming the activity and phosphorylated substrates of PKA, CAMKII, and other kinases as critical regulators. Time-resolved immunofluorescence of GSIS + GPCR signaling revealed previously unidentified PTMs accompanying β-cell organelle remodeling, including ciliary elongation, microtubule disassembly, Golgi remodeling, mitochondrial fusion and fission, and dynamic Golgi-mitochondrial contacts. Analysis of ATAT1 and HDAC6 revealed a mechanism of “opposing” phospho-regulated enzymes that control microtubule acetylation and, in turn, insulin secretion. Conserved phosphorylation and organelle responses in mouse and human islet cells support the relevance of these findings to human health. Together, our work provides unprecedented resolution of how metabolic and GPCR signals govern β cell signal transduction to tune insulin secretion. These data establish Metabolic STAMP as a predictive resource for studying the stimulus-dependent mechanisms underlying GSIS in β-cells.

Insulin secretion is regulated by multiple inputs to β cells, principally glucose, but also crucial physiological signals like metabolites and hormones. Findings from Metabolic STAMP here support that insulin secretion is tuned by compartmentalized and intersecting phosphorylation signaling inputs. Our analyses revealed that this signaling complexity is orchestrated through substrate specificity of kinase recognition motifs, spatial compartmentalization by scaffold proteins, engagement of distinct subcellular organelles, temporal signaling dynamics, as well as regulatory feedback. In contrast, agonists of GLP-1R and FFAR4 elicited partially overlapping yet distinct phosphorylation signatures not activated by glucose alone, including GLP1-R activation of ERK signaling and FFAR4 regulation of mitochondrial AKAP/PDE complexes and ciliary elongation. Thus, glucose establishes a basal framework for insulin secretion that is ’fine-tuned’ by GPCR pathways that regulate cell responses in time and in distinct subcellular compartments.

Metabolic STAMP also revealed hundreds of phosphorylation events *attenuated* by combining GLP1-R and FFAR4 agonists, indicating that regulatory crosstalk and feedback mechanisms could prevent excessive or maladaptive signaling in the β cell. For example, our studies revealed evidence of delayed phospho-relays and suppressive phosphatase activity stimulated by these GPCRs. Precedents for such feedback loops include GABAergic suppression to increase neural excitability, or hyperphosphorylation of Rb to drive cell cycle progression.^57–60^ Further STAMP studies with higher temporal resolution than achieved here could decipher the kinetics of this potential feedback loop to regulate the robustness and flexibility of β cell insulin secretion. Revealing a compartmentalized, stimulus-specific GPCR signaling architecture in β cells could explain how different pharmacological agents yield additive or synergistic effects on β cell function and suggest new combination therapeutics, tailored to modulate insulin secretion.

Though previous studies have explored the dynamics of context-specific fluctuations in phosphorylations during GSIS,^1, 2^ little was known about the spatial organization (if any) and function of these PTMs. Our study revealed transient molecular signals with localized changes in β cell organelle function during GSIS. Thus, different stimuli (glucose ± GPCR agonists) are transduced by common molecular machinery but in a temporal sequence and compartment-specific manner. For example, TUG891 upregulated mitochondrial AKAP1[S103] phosphorylation, while Ex4 stimulated ERK1[Y204] phosphorylation, a classic “ON” signal.^61^ Thus, our work suggests how combined stimulation of FFAR4 and GLP1-R during GSIS leads to enhanced insulin secretion by activating orthogonal but cooperating phosphotargets (**Fig. S1A**) Our GPCR studies also reveal unrecognized or little-characterized PTMs in modulating insulin secretion, including phosphorylation of ciliary proteins like KIF3A[S689], CEP170[S872], CEP97[S469] induced by FFAR4 (**Fig. 1D-E**), and phosphorylation of vesicular trafficking proteins like SEC22B[S137], MIA2[S1131], SNAP23[S110], and STXBP5[S759]) induced by Ex4 (**Fig. 1D-E**).

In addition to cilia, mitochondria, and cytoplasmic vesicles, Metabolic STAMP revealed dynamic PTMs in microtubules and Golgi, organelles previously linked to regulation by glucose but not GPCRs in β cells. Microtubules undergo dynamic rearrangement during insulin granule release,^45, 50, 62^ suggesting pivotal regulatory roles in GSIS. Our phosphoproteomics identified ATAT1[S315] and HDAC6[S21] as prominent cytoskeletal phosphosites (**Fig. 5A-B**), largely regulated by glucose (**Fig. 5C-F**). ATAT1 and HDAC6 are crucial enzymes with roles in reversible acetylation of α-tubulin^51^: phospho-ATAT1[S315] promotes α-tubulin[K40] acetylation and microtubule stabilization^52, 53^ , while phospho-HDAC6[S21] enhances deacetylase activity, driving microtubule deacetylation and depolymerization.^63^ Immunofluorescence and selective HDAC6 inhibitor studies here suggest that microtubule acetylation and deacetylation dynamics play an important role in GSIS, and GPCR stimulation helps to regulate these dynamics (**Fig. 6**). Our studies revealed differential localization of ATAT1, regulated by GPCR stimulation. ATAT1[S315] has previously been reported to induce nuclear exit, consistent with our data.^52^ Further studies are needed to establish the function of these PTMs during GSIS. Other than regulating microtubule stability to control vesicular traffic in β cells, we speculate that microtubule acetylation could also (or alternatively) increase the processivity of motor proteins, enabling enhanced kinesin and dynein traffic as seen in neurons.^64^ These data support a model in which ATAT1 and HDAC6 serve as a critical signaling ’node’ during GSIS to differentially regulate acetylated microtubule pools in a stimulus-dependent manner (**Fig. 6**).

How do FFAR4 and GLP1-R differentially regulate the postulated HDAC6/ATAT1 signaling node? We detected decreased HDAC6 or ATAT1 phosphorylation after FFAR4 or GLP-1R stimulation during GSIS, suggesting that kinase and/or phosphatase may dictate HDAC6/ATAT1 localization and function, which possibly control first and second-phase insulin secretion; these are testable possibilities for further work. With Metabolic STAMP, we also detected transient, stimulus-dependent contacts between *cis*-Golgi, mitochondria, and vesicular traffic machinery (**Fig. 4**). Interactions between mitochondrial and ER are better studied than mitochondrial-Golgi contacts;^65, 66^ the latter have been proposed to modulate membrane composition,^67^ calcium signaling, mitochondrial fission/fusion dynamics,^68^ and energetics.^65, 66^ Imaging showed GPCR-modulated mitochondrial-Golgi contacts and phosphorylations of mitochondria-Golgi contact regulatory proteins, including oxysterol-binding protein (OSBP).^69^ Further work is needed to decipher the function of these contacts during GSIS.

To strengthen our conclusions from studies of immortalized β cell lines here, we applied Metabolic STAMP to cadaveric human islets. Overall, the phosphoproteomics data from MIN6 studies recapitulated findings from human islet studies. Conversely, data from human islet imaging, phosphoproteomics, GWAS gene colocalization, and comparisons with prior Patch-seq analysis^70^ strongly supported our findings with MIN6 and MIN6-6 cells. For example, previous GWAS and Patch-seq studies of T1DM and T2DM revealed hundreds of risk loci, or protein-coding genes with disease risk. A number of these are found to be regulated by phosphorylation in MIN6 and/or human islets, pointing to possible mechanism for future study.^54, 55, 70^ Unexpectedly, loss of acetylated microtubules upon high glucose stimulation in primary human β cells was suppressed and delayed compared to MIN6 cells, suggesting the possibility of additional signaling complexity or lack of complete conservation in human β cells, whose regulation differs from rodent β cells (**Fig. 6A**). Other aspects of glucose-dependent microtubule deacetylation dynamics were conserved. Note that microtubule acetylation/de-acetylation patterns are not universal across the islet, as human islet α cells contain virtually no acetylated microtubules within conditions of our experiments, based upon our IHC studies. This suggests cell type specificity of cytoskeletal dynamics for different endocrine cell types, which would be an interesting future field of study to explore.

Our findings provide a resource to formulate hypotheses and heuristically frame future studies of signaling mechanisms regulating β cell GSIS and response to GPCRs. This includes studies to interrogate how differential GPCR stimulation leads to organelle-based regulation of PTMs in time and space to regulate GSIS. Further phosphoproteomic and imaging studies in primary human islets could identify singularities or similarities to islets from mice and other experimental models, including stem cell-derived human β cells. We foresee application of our approach to other important islet cell subsets, like α and δ cells. If so, our signaling and organelle roadmap should accelerate progress toward examining and validating candidate islet ’replacement’ cells proposed for use in autoimmune diabetes and other diseases.

## Limitations of the Study

The majority of these studies were performed in the mouse β stable cell lines MIN6 and MIN6-6 because of A) ability to expand to larger sample sizes for phosphoproteomics and imaging, B) the fact that it is a homogeneous cell population rather than a population of mixed cell types, thus increasing signal to noise in our time-resolved phosphoproteomics and imaging assays, and C) ease of access and culturing. Molecular mechanisms and timing of signaling events may not be entirely conserved between humans and mice, so future work should be done to dissect which pathways and processes are shared. Simultaneously, phosphoproteomics interpretations from human cadaveric islets are limited because of the mixed population of cell types. We chose not to dissociate islets into individual cell types to preserve cellular structure of as biologically relevant a model as possible. Our data already demonstrate cell-type specificity in cellular phenotypes during GSIS (*e.g.*, the lack of acetylated tubulin in α cells compared to β cells in the human islet (**Fig. 6, Fig. S6**)). As noted in **Fig. 6**, though, we observe substantially conserved phosphosignatures in response to stimuli between our human cadaveric islet and MIN6 samples, and previous studies have confirmed that MIN6 is an efficacious model of the GSIS response. This resource article serves as a foundation for much-needed research into the mechanisms underlying the function and regulation of GSIS in human β cells.

The sample sizes for human cadaveric islets were limited in this study due to limited tissue availability, both in access to donations and volume of islets. We also limited the scope of this study to adult, non-diabetic patient samples. Factors such as ethnicity, gender, age, BMI, environmental factors, etc. have not been fully considered in this study. Future work should be done to explore how such variables may influence signaling and cellular dynamics during GSIS in human patients and why there may be variability between samples.

Another limitation of these studies is the use of Exendin-4 as the agonist for GLP1-R rather than GLP-1, the endogenous ligand. In this study, we focused on Exendin-4 as our ligand of choice to achieve robust, sustained receptor engagement during GSIS and signaling assays. Exendin-4 is a high-affinity, long-acting GLP-1R agonist that is substantially more resistant to degradation than native GLP-1.^71–73^ The ability to maintain stable agonist levels over the course of our experiments was important for reducing variability due to peptide instability, therefore allowing us to achieve more robust phosphoproteomics and imaging data. We established this reference dataset as a resource for dissecting the differential signaling pathways regulating GSIS. Future work would benefit from exploring the degree to which the cellular and molecular phenotypes revealed in this study are consistent using endogenous GLP-1, as well as across different GLP-1 ligands.

## DECLARATION OF INTERESTS

The authors declare no competing interests.

## RESOURCE AVAILABILITY

### Lead contact

Further information and requests for resources and reagents should be directed to and will be fulfilled by the Lead Contact, Peter Jackson.

### Materials availability

All unique/stable reagents generated in this study are available from the Lead Contact with a completed Materials Transfer Agreement.

### Data and code availability

The published article includes all datasets generated during this study.

## ACKNOWLEDGEMENTS

Research reported in this publication was supported by the NIH. The content is solely the responsibility of the authors and does not necessarily represent the official views of the NIH.

MOA was supported by Stanford’s Translational Research and Applied Medicine Pilot Grant. RET was supported by Stanford Dean’s Fellowship, NIH F32 1F32GM142180-01A1, and 1 K99 GM154060-01. PKJ was supported by NIH grants 5R01GM114276, 2TR01GM121565, and 5UL1TR00108502. Work in Kim Lab was supported by NIH awards (R01 DK107507; R01 DK108817; U01 DK123743; R01 DK126482), the Breakthrough T1D Northern California Center of Excellence (to SKK and Dr. Q. Tang), and support from the Snyder and Leavitt families. Work done in the Knowles Lab was supported by NIH (P30 DK116074, R01 DK116750, R01 DK120565, R01 DK106236; R01 DK137889). We thank the Barrett, Hathaway, Killian, and Westly families for generous support. AR is supported by the Finnish Foundation for Cardiovascular Research, Diabetes Research Foundation, Emil Aaltonen Foundation, Ida Montin’s Foundation, Biomedicum Helsinki Foundation, Orion Research Foundation, and the Finnish Medical Foundation.

Human pancreatic islet processing and phenotyping were supported by Stanford Islet Research Core funded by NIH P30 DK116074 (to SKK). We thank the Nathanael Gray lab for the donation of the 240-kinase inhibitor screen featured in this paper. This work was also made possible by the Stanford Cell Sciences Imaging Facility, especially through the support of Kitty Lee and Jon Mulholland. We extend special thanks to Israel Larios and Roy Ng for instrumental technical support throughout the detailed experimental procedures. We thank Dr. Wenbiao Chen (Vanderbilt) for their donation of MIN6-6 cells.

## AUTHOR CONTRIBUTIONS

Conceptualization-MOA, RET, YH, SKK, PKJ; Methodology-MOA, RET, YH, AA, AR; LAX; Software-MOA, RET, MAM; Investigation-MOA, RET, YH, AA, LEL, MAM, LAX, AR; Resources-JWK, SKK, PKJ; Writing-MOA, RET, YH, AA SKK, PKJ; Supervision-SKK, PKJ; Funding Acquisition-MOA, RET, AA, SKK, PKJ

## SUPPLEMENTAL FIGURES

**Figure S1: Overview of Metabolic STAMP (<u>S</u>ynchronized <u>T</u>emporal-Spatial <u>A</u>nalysis via <u>M</u>icroscopy and <u>P</u>hosphoproteomics) to decipher GPCR-kinase signaling network governing insulin secretion.**

**(A)** Murine β cells or human islet cells were treated by high glucose alone or together with GPCR agonists to stimulate insulin secretion. During secretion, the following time-controlled samples were collected for Metabolic STAMP: cell media, cell lysates, live cells, and fixed cells.

**(B)** Cell medium was analyzed for hormone secretion levels like ELISA and/or bioluminescence based high-throughput assays.

**(C)** Cell lysates were processed for DDA-PASEF phosphoproteomics to map phosphorylation sites of GPCR signaling pathways. Putative kinases were nominated based on *in silico* analysis and then validated using kinase inhibitor assays.

**(D)** Paired live and fixed cell analysis via confocal microscopy were used to assess context specific changes in the following cellular compartments: microtubule cytoskeleton, mitochondria, Golgi and endosomal traffic, and primary cilia.

Figure was generated using BioRender.

**Figure S2: Mapping the differential phosphoproteomics of GSIS with and without co-stimulation with GPCR agonists (Expanded data from Figure 1**)

**(A)** Upset plot of overlapping phosphorylation events reduced in different GPCR stimulation conditions or glucose alone in MIN6 cells (>2-fold decrease, p<0.001).

**(B-F)** UMAP analysis reveals the differential effects of glucose and GPCR stimulation (FFAR4 vs GLP1-R) upon the cellular phosphoproteomic landscape. The size of the circle indicates log2 fold change, color of circle represents the known subcellular compartment where that protein functions. The first panel indicates the baseline of all the possible phosphorylations detected, while the subsequent panels show increased circle sizes for the phosphorylations detected that show marked changes in abundance in that stimulation condition.

(**G**) Schematic highlighting the key components of PKA signaling analyzed for differential phosphorylation in the right panels. Subcellular localization of the depicted AKAPs and PDEs is indicated below.

(**H-K**) Phosphorylation Change in PKA/AKAP/PDE Complex (Expanded data from Figure 1). Schematic of phosphorylation changes in individual components of PKA signaling. Phosphorylation abundance is represented as log2 fold change, with red indicating increases and blue indicating decreases.

(**L**) Glucose-induced CDK-motif phosphorylation on the cortical microtubule plus-end scaffold, reduced by GLP1-R and FFAR4 stimulation. Schematic highlighting the complex of plus-end microtubule regulating proteins that demonstrate the same pattern of phosphorylation in response to GSIS regulatory stimuli; each have a CDK consensus motif.

(**M**) Western blot of pERK1/2 validates findings made via mass spectrometry. Western blot for pERK1/2 (T202, Y204) phosphorylation confirming findings from mass spectrometry, with samples collected at 5min versus 20min post-stimulation. β-actin and Lamin are used here as loading controls.

(**N**) GSIS of healthy adult human cadaveric islet using perifusion assay to measure insulin secretion. This is the same batch of human islets used for performing phosphoproteomics. Concentration of insulin (mU/L) was measured at 15minutes, the same time point at which the human phosphoproteomics was performed. Error bars = SEM. Patient was non-diabetic, BMI=37, female, 18yo.

**Figure S3**: **Workflow for identification of key kinase regulators of GSIS and their substrates**. Schematic of the kinase inhibitor screen workflow for identifying novel candidate kinases regulating GSIS. MIN6-6 cells (which secrete luciferase-tagged insulin^67^) are seeded, serum-starved for 24hr to induce cell synchrony, and incubated in low glucose along with the Grey Lab Inhibitor Library, which consists of 240 different inhibitors at 50μM each. Cells were then switched to high glucose+GLP1-R+FFAR4 agonist treatment for 30 minutes. Samples were then assessed via NanoGlo Luciferase plate-reader assay before being binned based on the degree with which each kinase inhibitor increases/decreases GSIS compared to baseline. We can then focus on the functions of a specific kinase of interest by assessing phosphosite abundance in High Glucose + TUG891 + Exendin-4 treatment with and without supplementation with inhibitor treatment. All sites were scored based on kinome motifs and ranked based on fold change/p-value. The changes in site abundance sorted based on kinase motif was used as a tool for predicting the effects of different inhibitor treatments on different kinase activities.

**Figure S4:**

**(A)** Plot showing the dose-response effects of HDAC6i treatment on differential GSIS treatments in GSIS. Cells were preincubated with different doses of HDAC6i (with DMSO as control) for 15 min before stimulation with the following GSIS conditions: LG, HG, HG+TUG891, HG+Exendin4, and HG+TUG891+Exendin4 on GSIS. The following doses of HDAC6i were tested: 5nM, 50nM (IC_50_), 500nM. N=3

**(B)** Representative image showing ATAT1 co-localization with cis-Golgi (GM130) in MIN6 cells. Images were collected at 63x magnification; green= ATAT1, red= GM130, white= acetylated tubulin, blue= DAPI. Scale = 10µm.

**(C)** ATAT1 has differential cytoplasmic distribution upon stimulation with different GSIS conditions. Z-stack immunofluorescent images collected via confocal microscopy at 63x magnification at 5 versus 20min post-stimulation. The following markers were used for staining: ATAT1 (green), ꞵ-tubulin (red), acetylated tubulin (white), and DAPI (blue). Scale bar= 10μm. This is an expanded version of Figure 6H.

**Fig. S5: ATAT1 has differential localization to the nucleus upon stimulation with different GSIS conditions.** Z-stack immunofluorescent images collected via confocal microscopy at 63x magnification at 5min versus 20min post-stimulation. Images show the following markers for localization: ATAT1 (green) and DAPI (blue). Scale bar= 10μm.

**Fig. S6: Glucagon-positive α cells in islets from human patient donors have little to no acetylated tubulin across conditions.** Confocal microscopy (60x magnification, oil, z-stacks with 0.28µm per slice) was performed of each experimental condition. Representative images are shown here of the 15min time point for LG. DAPI= blue, glucagon= green, insulin= red, acetylated tubulin= white. Scale bar = 10μm for 60x image. Images were processed using FIJI imaging software.

## METHODS

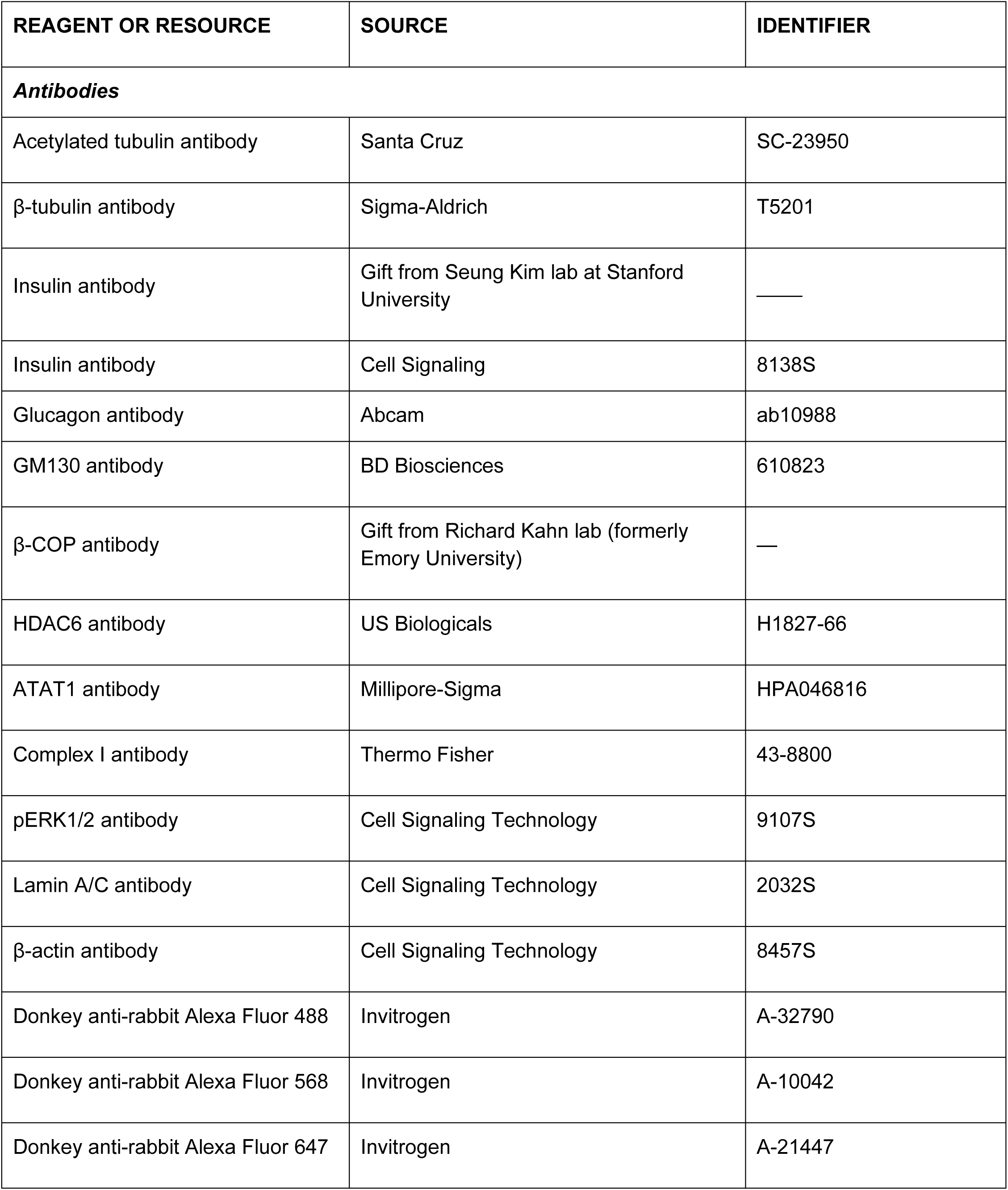
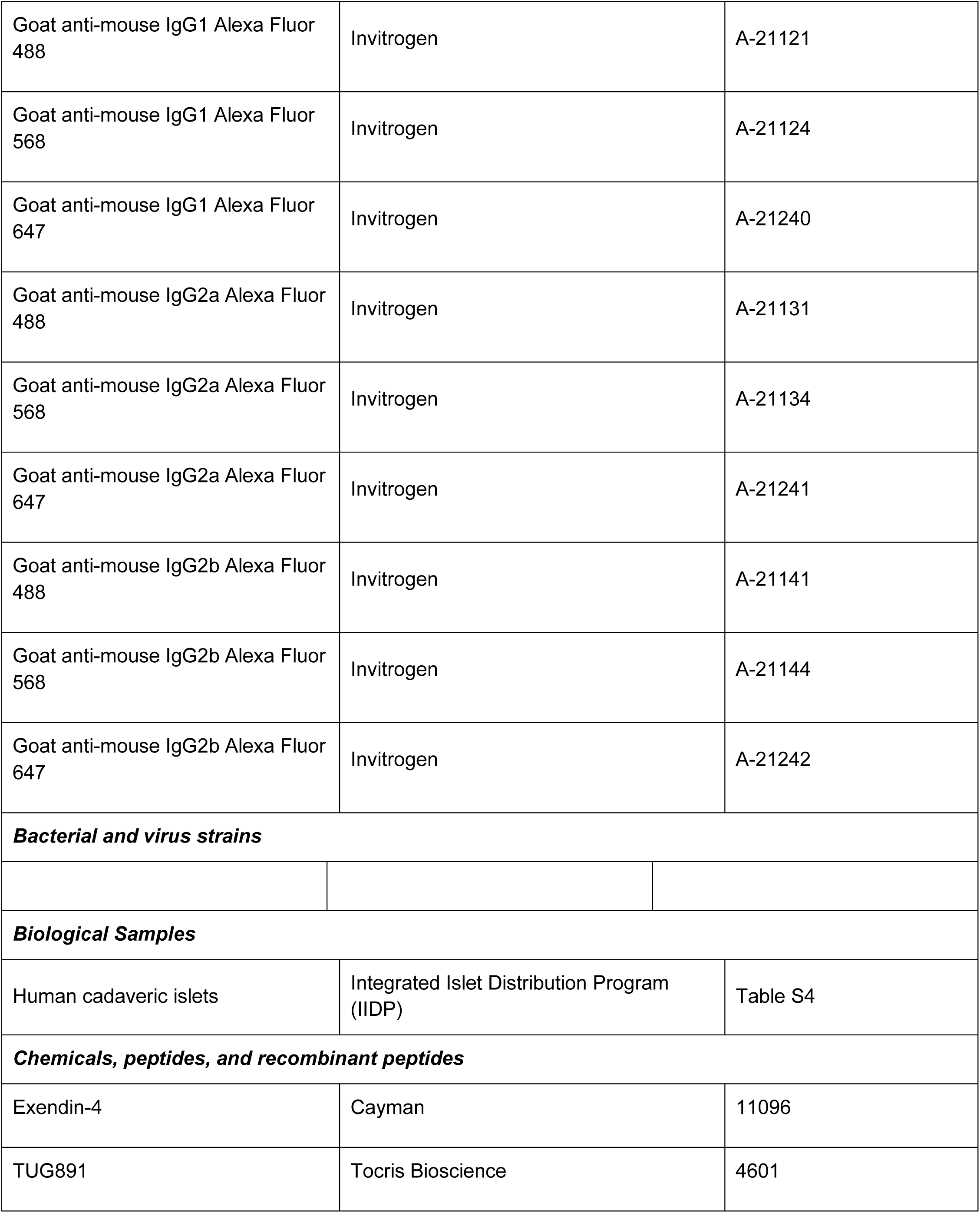
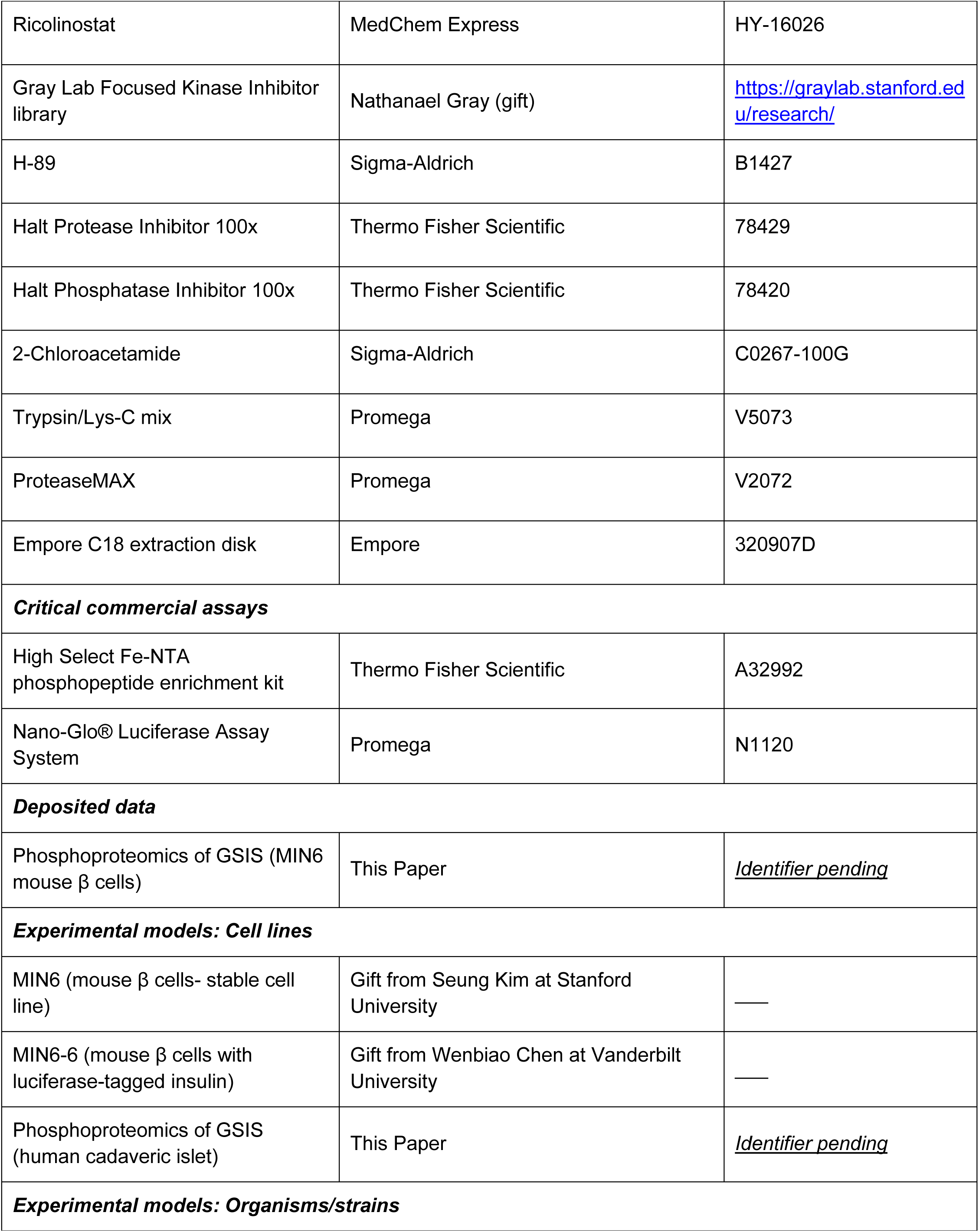
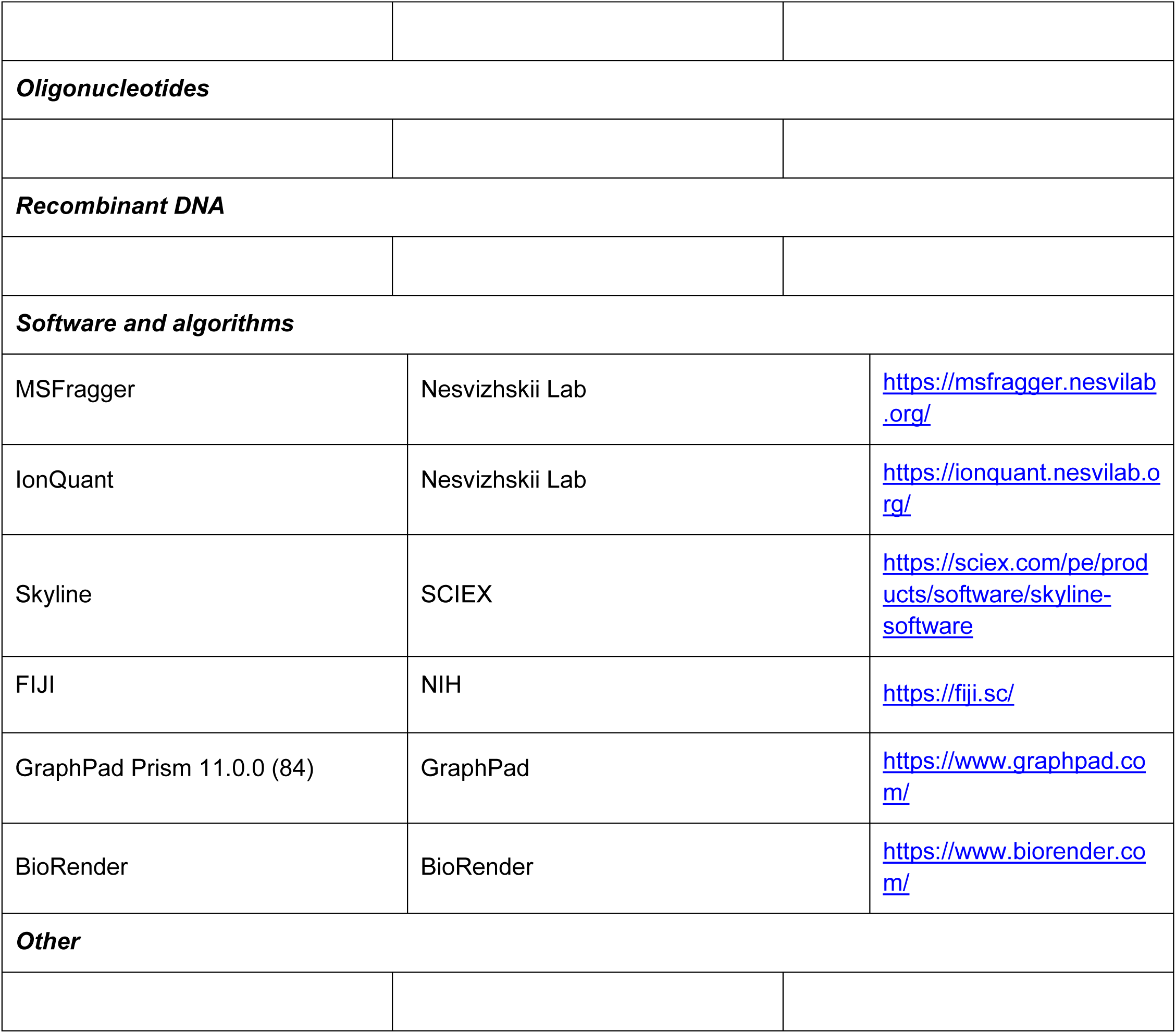
KEY RESOURCES TABLE.

## EXPERIMENTAL MODEL AND STUDY PARTICIPANT DETAILS

### MIN6 and MIN6-6 Mouse β cells

Mouse pancreatic MIN6 β cells were gifted from Seung Kim’s lab at Stanford University. Pancreatic MIN6-6 cells were gifted from Wenbiao Chen’s lab at Vanderbilt University. Stable cell lines were cultured in DMEM medium (buffered with sodium bicarbonate (Gibco; 25080-094) containing 10% heat-inactivated fetal bovine serum (Gemini; Cat #: 100-106), 10mM HEPES (Gibco; Cat #: 15630080) 0.1% v/v β-mercaptoethanol (Invitrogen; Cat #: 21985023), and 1% pen/strep (Gibco; Cat #: 10378016). Cells were maintained at low passage (below passage 30) to avoid genetic drift and checked at least monthly for mycoplasma using Hoechst staining and immunofluorescence. For all imaging and mass spectrometry experiments, 5 × 10×6 cells were plated into 10cm dishes. For all insulin secretion assays, 1 × 10 ×5 cells were plated per well of a 96-well dish. MIN6 cells were cultured and subjected to serum starvation in 5 mM glucose for 24 hours prior to stimulation. After preincubation with low glucose (2.7 mM) for one hour, cells were stimulated under the following conditions (each compared with low glucose control): High Glucose (HG: 17.8mM), HG + 20nM Ex4, HG + 100µM TUG891, HG + 20nM Exendin-4 + 100µM TUG891 (Combo), Combo + 10µM NKY-80 (ADCY inhibitor), Combo + 1µM H-89 (PKA inhibitor), Combo + 1µM KN-93 (CAMKII inhibitor). For the phosphoproteomics cells are lysed at 10 minutes. For the imaging cells stimulated for durations of 5 and 20 minutes as indicated, followed by sample harvest for downstream analysis.

### Human Cadaveric Islets

Human cadaveric islets were obtained through Stanford Diabetes Research Center’s Islet Research Core. These samples were harvested from de-identified human donors. These samples were harvested less than 15hr of cold ischemia time. Islets were spun down at 200 g (2-4 Minutes) to remove the culture medium, washed with ice-cold 1x PBS, and spun down again at 200 rcf. Samples were then incubated with 190μL 2.8mM Low Glucose for two hours at 37°C before use for future assays (described below). For mass spectrometry experiments, 3000 IEQ were plated for each condition. To induce stimulated GSIS in isolated human pancreatic islets, cells were treated with 310μL of the following conditions: 2.8 mM glucose (LG), 25mM glucose alone (HG), 25mM glucose with the FFAR4 agonist TUG891 (161μM), or 25 mM glucose with Ex4 (32.2nM). Following drug addition, cells were incubated at 37°C for 15min. After incubation, the culture medium was collected for subsequent ELISA. Cells were then lysed by addition of 300μL lysis buffer and immediately boiled for 5 minutes at 95°C. Lysates were stored at −80°C until further preparation for phosphoproteomic analysis.

## METHOD DETAILS

### Reagents, antibodies, and plasmids

The following primary antibodies were used in this study. **<u>For IF</u>**: acetylated tubulin (mouse IgG2B; Santa Cruz; Cat #: SC-23950; 1:2000 of 200 μg/ml stock), ꞵ-tubulin (mouse IgG1; Sigma-Aldrich; Cat #: T5201; 1:2000 of 2mg/mL stock)), insulin (guinea pig; discontinued antibody gifted from Seung Kim Lab;^68,69^; 1:250 of stock), GM130 (mouse IgG1; BD Biosciences; Cat #: 610823; 1:500 of 250µg/mL), ꞵ-COP (rabbit; gifted from Richard Kahn lab;^70^ 1:1000 of stock), HDAC6 (Rabbit; US Biologicals; Cat #: H1827-66; 1:250 of 0.1mg/mL stock), ATAT1 (Rabbit; Millipore-Sigma; Cat #: HPA046816; 1:250 dilution of 0.1mg/mL stock), Complex I (mouse Ig2B; Thermofisher; Cat #:43-8800; 1:1000 of 1mg/mL stock). **<u>For Western blot</u>**: pERK 1/2 (mouse IgG1; Cell Signaling Technologies; Cat #: 9107S; 1:2000 dilution), Lamin (rabbit IgG1; Cell Signaling; Cat #:2032S; 1:1000 dilution of 0.5mg/mL stock), and ꞵ-actin rabbit IgG; Cell Signaling; Cat #: 8457S; 1:2000 of 1mg/mL stock).

The following Alexa Fluor secondary antibodies (Invitrogen) were used in this study: (i) Donkey-anti-Rabbit 488 (Cat #: A-32790), (ii) Donkey-anti-Rabbit 568 (Cat #: A-10042), (iii) Donkey-anti-Rabbit 647 (Cat #: A-21447), (iv) Goat-anti-mouse IgG1 488 (Cat #: A-21121), (v) Goat-anti-mouse IgG1 568 (catalog no. A-21124), (vi) Goat-anti-mouse IgG1 647 (Cat #: A-21240), (vii) Goat-anti-mouse IgG2a 488 (Cat #: A-21131), (viii) Goat-anti-mouse IgG2a 568 (Cat #: A-21134), (ix) Goat-anti-mouse IgG2a 647 (Cat #: A-21241), (x) Goat-anti-mouse IgG2b 488 (Cat #: A-21141), (xi) Goat-anti-mouse IgG2b 568 (Cat #: A-21144), (xii) Goat-anti-mouse IgG2b 647 (Cat #: A-21242)

The following drugs were used in this study: Exendin-4 (Cayman; Cat #: 11096), TUG891 (Tocris Bioscience; Cat #: 4601), Ricolinostat (HDAC6 inhibitor) (MedChem Express; Cat #: HY-16026), H-89 (PKA inhibitor) (Sigma-Aldrich; Cat #: B1427). Sterile DI water or DMSO (Sigma-Aldrich; Cat #: 276855) was used for diluting drugs. The kinase inhibitor screen (240 inhibitors tested) was shared with us by the Nathanael Grey lab^32–34^ in 96-well plate format (https://graylab.stanford.edu/probe-resources/) with the Stanford High-Throughput Screening Core (Stanford HTS).

The following reagents are used for phosphoproteomics: Halt™ Protease Inhibitor 100x (Thermofisher Scientific; Cat #: 78429), Halt™ Phosphatase Inhibitor 100x (Thermofisher Scientific; Cat #: 78420), 2-Chloroacetamide (CAA) (Sigma-Aldrich; Cat #: C0267-100G), HEPES, >99.5% Titration (Sigma-Aldrich; Cat #: H3375-5KG), Blunt-End Needles, 16 Gauge (STEMCELL; Cat #: 28110), Acetonitrile (ACN) (Honeywell; Cat #: 14261-1L), Sodium Chloride (NaCl) (Cat #: S3014-5KG, Sigma-Aldrich), Methanol (Millipore Sigma; Catalog #: 900688-1L), Water, Optima™ LC/MS Grade (Fisher Chemical; Cat#: W6-4), Trypsin/Lys-C Mix, Mass Spec Grade (Cat #: V5073, Promega), ProteaseMAX (Promega; Cat #: V2072), Ammonium Bicarbonate (ABC) (Honeywell; Cat #: 40876-50G), Pierce™ Premium Grade TCEP-HCl (Thermofisher Scientific; Cat #: PG82080), Pierce™ BCA Protein Assay Kit (Thermofisher Scientific; Cat #: 23225), Pierce™ Bovine Serum Albumin Standard Pre-Diluted Set (Thermofisher Scientific; Cat #: 23208), Pierce™ Quantitative Colorimetric Peptide Assay (Thermofisher Scientific; Cat #: 23275), High Select Fe-NTA Phosphopeptide Enrichment Kit (Thermofisher Scientific; Cat #: A32992), Empore C18 47mm Extraction Disk, Model 2215 (Empore; Cat #: 320907D), Formic acid, 99.5%, Optima™ LC/MS Grade (Thermofisher Scientific; Cat #: A117-50), Trifluoroacetic acid (TFA) (Fisher Scientific; Cat #: AAL06374AC)

### Kinase Inhibitor Screen on Glucose-Stimulated Insulin Secretion (GSIS) in MIN6 cells (Table S3, Fig.S2)

MIN6-6 cells were first cultured under standard conditions and then seeded into 96 multiwell plates at 80% confluency for 24 hours to allow attachment and growth. After initial culture, cells were serum-starved for 24 hours at 5 mM Glucose to synchronize metabolic activity. To initiate the kinase inhibitor screen, cells were pre-incubated for 1hr in low glucose (2.8mM) medium with 5 µM of each compound from the Gray Lab kinase inhibitor library (240 drugs). Following this pre-treatment, cells were stimulated for 30min at high glucose (17.8 mM) in the continued presence of the kinase inhibitors, and 100µM TUG891 and 20nM Ex4, including low glucose and high glucose controls. After stimulation, glucose-stimulated insulin secretion was quantified using the Nano-Glo® Luciferase Assay according to the manufacturer’s instructions, with luminescence measured to assess GSIS across conditions.

### Phosphoproteomics of MIN6 and Human Cadaveric Islets

Stock solutions were prepared, including 1 M tris-HCl (pH 8.5) and 5 M potassium hydroxide (KOH). Plates (10cm) containing 8 × 10×6 MIN6 cells were harvested for each replicate and time point for phosphoproteomic analysis. SDC lysis buffer was prepared fresh using 4% (w/v) SDC, 100mM tris-HCl (pH 8.5), and 1× Halt protease and phosphatase inhibitor cocktail. The buffer was made fresh. A 1ml volume of lysis buffer was added to each plate. Lysates were immediately heat-treated for 5min at 95°C to facilitate lysis and inactivate endogenous proteases and phosphatases. Then, lysates were homogenized by sonication at 4°C.

Reduction/alkylation buffer included 100mM TCEP and 400mM CAM, pH 7-8 adjusted with KOH, prepared immediately before use to preserve CAM activity. Then, disulfide bonds and carbamidomethylated cysteine residues were reduced by adding a 1:10 volume of reduction/alkylation buffer to the samples. Samples were incubated for 30min at room temperature at 1000 rpm, in the dark. After removing the samples from the shaker, Lys-C and trypsin enzymes were added at a 1/100 ratio of protein/enzyme ratios, and the samples were digested for 16hrs at 37°C with shaking at 1000 rpm.

Following overnight digestion, peptides were acidified with TFA, centrifuged, and cleaned up using Sep-Pak tC18 1 cc columns. The purified peptides were then quantified using the ThermoFisher Pierce™ Quantitative Peptide Assay to ensure accurate input amounts for downstream enrichment. Subsequently, phosphopeptides were enriched using the High Select Fe-NTA IMAC phosphopeptide enrichment kit (ThermoFisher) following the supplier’s instructions for binding, washing, and elution steps.

Homemade Stage-Tips were constructed using two C18 Empore disks, following established procedures. The fabricated Stage-Tips were washed two times with 100μL of methanol, one time with 100μL of 80% acetonitrile/0.1% acetic acid, and two times with 100μL of 1% acetic acid. Enriched phosphopeptides were loaded onto the Stage-Tips in 100μL of 1% acetic acid. Subsequently, the Stage-Tips were washed three times with 100μL of 1% acetic acid to remove salts. Last, the phosphopeptides were eluted from the Stage-Tips using two elution steps of 30μL each, with 80% acetonitrile/0.1% acetic acid as the elution buffer.

### Liquid chromatography setup

A nanoELute ultrahigh-pressure nanoflow chromatography system was used and directly coupled online with a hybrid trapped ion mobility spectrometry-quadrupole time-of-flight mass spectrometer (timsTOF HT, Bruker) using a nanoelectrospray ion source (CaptiveSpray, Bruker Daltonics).

### Chromatographic conditions

The liquid chromatography was conducted at a constant temperature of 50°C, using a reversed-phase column (PepSep column, 10 cm by 150 μm ID, packed with 1.5μm C18-coated porous silica beads, Bruker) connected to the 10μm emitter (Bruker). The mobile phase consisted of two components: Mobile Phase A, comprising water with 0.1% acetic acid (v/v), and Mobile Phase B, comprising ACN with 0.1% acetic acid (v/v).

### Gradient elution

Peptide separation was achieved using a linear gradient from 2 to 33% Mobile Phase B within 60 min. This was followed by a washing step with 95% Mobile Phase B and subsequent re-equilibration. The chromatographic process maintained the flow rate at 400nL/min.

### MS acquisition

Samples were analyzed using the timsTOF HT mass spectrometer in DDA-PASEF mode. The TIMS elution voltage was calibrated linearly to obtain reduced ion mobility coefficients (1/K0) by using three selected ions from the Agilent ESI-L Tuning Mix [mass/charge ratio (m/z) 622, 922, and 1222]. The mass and ion mobility ranges were set from 100 to 1700 m/z and 0.7 to 1.3 1/K0, respectively. Both ramp and acquisition times were set at 100ms. Precursor ions suitable for PASEF-MS/MS were chosen from TIMS-MS survey scans using the PASEF scheduling algorithm. A polygon filter was applied to the m/z and ion mobility plane to prioritize features likely representing peptide precursors over singly charged background ions. The quadrupole isolation width was set to 2 Th for m/z < 700 and 3 Th for m/z > 700, with collision energy linearly increased from 20 to 60 eV as ion mobility ranged from 0.6 to 1.6 (1/K0).

### Mass Spectrometry Data Analysis

Raw data files were processed using MS Fragger software against the NCBI Homo sapiens RefSeq protein database. Search parameters included CID (collision-induced dissociation) fragmentation with a precursor error tolerance of 10 parts per million (ppm) and a fragment ion tolerance of 20 ppm. Searches included S/T/Y phosphorylation and up to three modifications per peptide besides standard modifications. Peptides were validated using Percolator and Protein Prophet at 1% FDR (false discovery rate). Protein quantification was performed using IonQuant, with normalization across runs and Match Between Runs settings accommodating retention time and ion mobility tolerances of 0.4 min and 0.05 (1/K0), respectively.^71^

Using the comprehensive kinase substrate specificity profiles established by Cantley’s lab,^74^ we assigned the putative kinases for each phosphorylation site based on the tool available on PhosphoSite. The final kinase rankings were determined by calculating the percentile position of each kinase’s score within the distribution of scores generated from analyzing all available serine/threonine phosphorylation sites from PhosphoSite.^24^ The phosphorylation signaling maps are only plotted for the highly confident phosphorylations sites, manually inspected by Skyline. MS2 scans are extracted and confirmed for each phosphorylation site demarcated in the figures.

### UMAP Analysis Based on Localization with COMPARTMENT Database Annotation

All data processing and visualization were performed in Python (v3.11) using pandas, seaborn, and matplotlib, with an interactive interface built in Streamlit to facilitate dynamic data exploration and visualization.

Experimental phosphoproteomic data tables were analyzed with FragPipe-Analyst for the log₂ fold change values and associated p-values. A background protein list containing all phosphoproteins served as the reference set for generating the UMAP embedding. This background enabled a standard coordinate space onto which experimental data were overlaid. Quantitative phosphoproteomics data were merged into a unified kinase-substrate-level table, consolidating all condition measurements per protein in a structured format. The COMPARTMENT database^72^ was employed to provide subcellular localization annotations for all proteins in the background protein list. This database integrates evidence from diverse sources, including experimental data, computational predictions, and literature text mining, to assign proteins to specific cellular compartments. These localization annotations enriched the biological interpretation of the UMAP embedding by linking proteomic changes to their respective subcellular contexts. Uniform Manifold Approximation and Projection (UMAP) was applied to the full background dataset to embed proteins in a two-dimensional space, preserving both local and global relationships among proteins. Condition-specific log₂ fold change and p-value data were mapped onto the background UMAP embedding. Visualization outputs consisted of: Condition-specific protein data visualized on the UMAP coordinate space.

### Human Islet Secretion Assay and Hormone Assessment

The human islets were recovered for 2 hours in RPMI media (Thermo Fisher) supplemented 2.8 mM glucose (Sigma-Aldrich), 1mM Sodium Pyruvate (Thermo Fisher), 10mM HEPES (Thermo Fisher), and 0.1% (w/v) BSA (Sigma-Aldrich) prior to static insulin secretion analysis. Batches of 3000 IEQs were then treated with glucose or GPCR agonists as indicated for 15 min at 37 degrees. Supernatants were collected at the end of incubation and insulin levels were quantified by human insulin ELISA kits (Mercodia).

### Cryosectioning of Human Cadaveric Pancreatic Islets

After stimulation of different GSIS conditions, cells were fixed for 15 minutes with 4% PFA/PBS at 37°C. Samples were then spun down and washed 2x with PBS, spinning down and removing the supernatant from each wash. Using the Collagen Gel Cell Culturing Kit (FUJIFILM Wako; Cat #:638-00781), make a master mix of collagen components A and B (v: v = 8:1) enough for all tubes (usually 300μL per tube). Mix completely by vortexing. *It is recommended to use more AB mix than necessary at this point and remove the extra later.* Aliquot the AB mix to islet pellet and mix very well. Then, quick spin to collect islets at the bottom of the tube. Add component C (A: B:C = 8:1:1) to each tube, mix immediately by vortexing and then quick spin to sink islets to bottom. Collagen solidifies quickly at room temperature upon the addition of component C, so make sure to keep the samples on ice at all idle times. Incubate samples on ice for 10 minutes to allow the islets to immerse in the solution completely. Afterward, incubate the mix in a 37°C waterbath to allow the samples to solidify, ∼30min. Use a pipet tip to poke the top of collagen and swirl gently to check whether the samples have solidified with the islets locked in place. Add 4% PFA/PBS to further fix for 30min-1hr at 4°C. Afterward, take off PFA to stop fixation and wash with cold PBS twice.

To prepare the samples for cryosectioning, pour the collagen chunk out (or pull out gently with tweezers) onto a petri dish. Cut off the extra collagen above the islet cluster. Then, put the small collagen piece back into the Eppendorf tube and add in 30% sucrose. Incubate the samples overnight at 4°C or until the sample has sunk to the bottom of the tube.

After ensuring that the sample has sunk to the bottom of the tube, carefully take out the sample using tweezers onto a petri dish. Trim the collagen block using tweezers under a dissecting microscope to avoid losing sample. The purpose is to make the sample as small and flat as possible before embedding. Use a p10 pipette to remove residual liquid completely to avoid ice crystal formation during cryopreservation.

Use Sharpie to draw a line around the collagen piece to mark the location of the sample. Then, add a small amount of OCT to just cover the collagen. Let the block sit a few minutes to allow the OCT to infiltrate into the collagen. Then, put it on dry ice. Once the OCT just starts turning white, add more OCT to the top of the mold and wait until the entire block solidifies. Store at -80°C until prepared to start sectioning.

Samples were prepared for immunohistochemistry by making 6µm cryosections. Samples were stored at - 80°C until prepared for immunohistochemistry.

### Fixed Cell Imaging of MIN6 Cells and Human Cadaveric Pancreatic Islets

MIN6 cells were prepared for immunofluorescence according to standard lab protocols.^13^ In brief, cells were previously plated onto glass coverslips (12CIR-1.5 (Fisherbrand; Cat #: 12-545-81), after being passed through a cell strainer (Fisher Scientific; Cat #: 22-363-548) to ensure plating of single cells and to reduce cell clumping. This allows for better resolution of subcellular compartments via microscopy. After growing the cells and performing the different experimental conditions, cells were fixed at 37°C with 4% PFA/PBS (diluted from 16% stock; Electron Microscopy Sciences; Cat #: 100503-917) for 15 minutes. Coverslips were washed with PBS 4x before being permeabilized for 10 minutes with 0.1% TritonX-100 (Fisher Scientific; Cat #: AC21568-2500). Samples were incubated in block buffer (1% BSA/PBS) (Sigma; Cat #: A8806) for 1hr at room temperature and then incubated with primary antibody diluted in blocking buffer overnight at 4°C (see above for specific dilutions for different primary antibodies). Samples were then washed 4x with PBS and incubated with secondary antibody diluted 1:500 in blocking buffer for 1hr at room temperature in the dark. Coverslips were washed 2x with PBS and then incubated in DAPI (4′,6-Diamidino-2-phenylindole (DAPI), 10 mg/mL: (Biotium; catalog no. 40043)) diluted in PBS for 5 minutes at room temperature in the dark. Samples were then washed 2x with PBS before being mounted onto slides (VistaVision Microscope Slides, Histobond (75 mm by 25 mm by 1mm): (VWR; catalog no. 16005-110)) with SlowFade Gold Antifade reagent: (Invitrogen; Cat #: S36936).

For immunohistochemistry of human cadaveric islet cryosections, frozen slides were thawed for at least 30 minutes at room temperature before use. Samples were then prepared for immunofluorescence using a similar protocol as above, except that samples were incubated in secondary antibody for 2hrs rather than 1hr.

### Western Blot

Post-treatment (5×10×6 cells) samples were collected and lysed in RIPA buffer, and clarified lysates were treated with an SDS loading buffer containing β-mercaptoethanol. Upon electrophoretic separation, Western blots were prepared using the eBlot L1 (Genscript) following the manufacturer’s protocols onto 0.2μm PVDF membranes (Thermo Fischer, Cat #: 22860). Post-transfer, membranes were blocked in Li-Cor Intercept® (TBS) Blocking Buffer (Cat #: 927-60001) overnight at 4°C and cut with razor blade at appropriate molecular weight markers. Appropriate membrane slices were incubated with the following antibodies diluted in blocking buffer for 1hr at room temperature: (Anti-mouse Acetylated Tubulin 1:2000; Cat#: sc-23950), (Anti-Rabbit phosphor-pERK 1:1000; Cat #: 9107S), (Anti-Rabbit Lamin A+C 1:1000, Cell Signaling Cat#: 2032), (Anti-Rabbit Beta-Actin; 1:1000; Cat #: A1978). Membranes were washed 5x in TBST and incubated with Li-Cor secondary antibodies diluted in blocking buffer (Donkey anti-Rabbit IRDye 800 1:15000 and Donkey anti-mouse IRDye 680 1:15000). After 30 minutes of incubation at room temperature, membranes were washed in TBST and imaged on the LICOR Odyssey Imager.

### Image Processing/Quantification

Image analysis was performed using FIJI imaging software, using different strategies based on the assay. For each experiment, all test groups were treated the same for image processing: z-stacking, subtracting background (along with Math-Subtract function after measuring background), adjusting brightness/contrast to baseline, cropping, etc. Representative images with scale bars are included for each experiment.

Ciliary length was assessed via manual measurements using the FIJI measuring tool.

The average volume of an individual mitochondrion for each condition was assessed using FIJI plugin 3D Mitochondrial Analyzer.^73^

Volume of microtubule density in the cell was measured by thresholding and creating a mask for the total cell area versus thresholding and creating a mask for the microtubule network. Default settings were used for thresholding, with manual inspection for each image to ensure that the mask corresponded well with the image. The area of each was measured, and the results were reported as the percentage of the total cell area that contained microtubules. Note that this is a measure of total microtubule density, not considering other features of microtubules like length, polymerized versus depolymerized pools, etc. A similar strategy was used for measuring acetylated microtubule density except that we measured the total percentage of microtubules that are acetylated (aka, =area of acetylated microtubules/area of total microtubules).

Cytoplasmic distribution of β-COP was measured, after processing all images using the same parameters to subtract background, by generating ROIs for the cytoplasm of each cell and measuring the Mean Gray Value.

For all other quantification, yes-no binning was used to calculate the percentage of cells with the given phenotype, ∼100 cells per replicate.

## QUANTIFICATION AND STATISTICAL ANALYSIS

Data analysis and plotting was performed using Microsoft Excel and GraphPad Prism. One-Way ANOVA was used for statistical analysis, with p values = \**P* = 0.05; \*\**P* = 0.01; ***P=0.001; \*\*\*\**P* = 0.0001. All experiments were performed in biological replicate. All image quantification was performed via FIJI imaging software. Final figures were prepared in BioRender.

## Notes

### Competing Interest Statement

The authors have declared no competing interest.

### Summary of Updates

We have revised the article to incorporate new data concerning human cadaveric islets, updated data analysis, and revised the order and formatting of figures. Furthermore, we revised the text to reflect these changes.

## REFERENCES

1. Sacco F, Humphrey SJ, Cox J, Mischnik M, Schulte A, Klabunde T, Schafer M, Mann M. Glucose-regulated and drug-perturbed phosphoproteome reveals molecular mechanisms controlling insulin secretion. Nat Commun. 2016;7:13250. Epub 20161114. doi: 10.1038/ncomms13250. PubMed PMID: 27841257; PMCID: PMC5114537.

2. Sacco F, Seelig A, Humphrey SJ, Krahmer N, Volta F, Reggio A, Marchetti P, Gerdes J, Mann M. Phosphoproteomics Reveals the GSK3-PDX1 Axis as a Key Pathogenic Signaling Node in Diabetic Islets. Cell Metab. 2019;29(6):1422–32 e3. Epub 20190314. doi: 10.1016/j.cmet.2019.02.012. PubMed PMID: 30879985.

3. Kulaj K, Harger A, Bauer M, Caliskan OS, Gupta TK, Chiang DM, Milbank E, Reber J, Karlas A, Kotzbeck P, Sailer DN, Volta F, Lutter D, Prakash S, Merl-Pham J, Ntziachristos V, Hauner H, Pfaffl MW, Tschop MH, Muller TD, Hauck SM, Engel BD, Gerdes JM, Pfluger PT, Krahmer N, Stemmer K. Adipocyte-derived extracellular vesicles increase insulin secretion through transport of insulinotropic protein cargo. Nat Commun. 2023;14(1):709. Epub 20230209. doi: 10.1038/s41467-023-36148-1. PubMed PMID: 36759608; PMCID: PMC9911726.

4. Li J, Li Q, Tang J, Xia F, Wu J, Zeng R. Quantitative Phosphoproteomics Revealed Glucose-Stimulated Responses of Islet Associated with Insulin Secretion. J Proteome Res. 2015;14(11):4635–46. Epub 20151015. doi: 10.1021/acs.jproteome.5b00507. PubMed PMID: 26437020.

5. Tang JS, Li QR, Li JM, Wu JR, Zeng R. Systematic Synergy of Glucose and GLP-1 to Stimulate Insulin Secretion Revealed by Quantitative Phosphoproteomics. Sci Rep. 2017;7(1):1018. Epub 20170421. doi: 10.1038/s41598-017-00841-1. PubMed PMID: 28432305; PMCID: PMC5430885.

6. Wu CT, Hilgendorf KI, Bevacqua RJ, Hang Y, Demeter J, Kim SK, Jackson PK. Discovery of ciliary G protein-coupled receptors regulating pancreatic islet insulin and glucagon secretion. Genes Dev. 2021;35(17-18):1243–55. Epub 20210812. doi: 10.1101/gad.348261.121. PubMed PMID: 34385262; PMCID: PMC8415323.

7. Friedman JM. The discovery and development of GLP-1 based drugs that have revolutionized the treatment of obesity. Proc Natl Acad Sci U S A. 2024;121(39):e2415550121. Epub 20240919. doi: 10.1073/pnas.2415550121. PubMed PMID: 39297680; PMCID: PMC11441540.

8. Meloni AR, DeYoung MB, Lowe C, Parkes DG. GLP-1 receptor activated insulin secretion from pancreatic beta-cells: mechanism and glucose dependence. Diabetes Obes Metab. 2013;15(1):15–27. Epub 20120801. doi: 10.1111/j.1463-1326.2012.01663.x. PubMed PMID: 22776039; PMCID: PMC3556522.

9. Kreymann B, Williams G, Ghatei MA, Bloom SR. Glucagon-like peptide-1 7-36: a physiological incretin in man. Lancet. 1987;2(8571):1300–4. doi: 10.1016/s0140-6736(87)91194-9. PubMed PMID: 2890903.

10. Hughes JW, Cho JH, Conway HE, DiGruccio MR, Ng XW, Roseman HF, Abreu D, Urano F, Piston DW. Primary cilia control glucose homeostasis via islet paracrine interactions. Proc Natl Acad Sci U S A. 2020;117(16):8912–23. Epub 20200406. doi: 10.1073/pnas.2001936117. PubMed PMID: 32253320; PMCID: PMC7184063.

11. Newcombe EA, Delaforge E, Hartmann-Petersen R, Skriver K, Kragelund BB. How phosphorylation impacts intrinsically disordered proteins and their function. Essays Biochem. 2022;66(7):901–13. doi: 10.1042/EBC20220060. PubMed PMID: 36350035; PMCID: PMC9760426.

12. Needham EJ, Parker BL, Burykin T, James DE, Humphrey SJ. Illuminating the dark phosphoproteome. Sci Signal. 2019;12(565). Epub 20190122. doi: 10.1126/scisignal.aau8645. PubMed PMID: 30670635.

13. Azizzanjani MO, Turn RE, Asthana A, Linde-Garelli KY, Xu LA, Labrie LE, Mobedi M, Jackson PK. Synchronized temporal-spatial analysis via microscopy and phosphoproteomics (STAMP) of quiescence. Sci Adv. 2025;11(17):eadt9712. Epub 20250425. doi: 10.1126/sciadv.adt9712. PubMed PMID: 40279433; PMCID: PMC12024681.

14. Johnson JL, Yaron TM, Huntsman EM, Kerelsky A, Song J, Regev A, Lin TY, Liberatore K, Cizin DM, Cohen BM, Vasan N, Ma Y, Krismer K, Robles JT, van de Kooij B, van Vlimmeren AE, Andree-Busch N, Kaufer NF, Dorovkov MV, Ryazanov AG, Takagi Y, Kastenhuber ER, Goncalves MD, Hopkins BD, Elemento O, Taatjes DJ, Maucuer A, Yamashita A, Degterev A, Uduman M, Lu J, Landry SD, Zhang B, Cossentino I, Linding R, Blenis J, Hornbeck PV, Turk BE, Yaffe MB, Cantley LC. An atlas of substrate specificities for the human serine/threonine kinome. Nature. 2023;613(7945):759–66. Epub 20230111. doi: 10.1038/s41586-022-05575-3. PubMed PMID: 36631611; PMCID: PMC9876800.

15. Varney MJ, Benovic JL. The Role of G Protein-Coupled Receptors and Receptor Kinases in Pancreatic beta-Cell Function and Diabetes. Pharmacol Rev. 2024;76(2):267–99. Epub 20240213. doi: 10.1124/pharmrev.123.001015. PubMed PMID: 38351071; PMCID: PMC10877731.

16. Dalle S, Abderrahmani A. Receptors and Signaling Pathways Controlling Beta-Cell Function and Survival as Targets for Anti-Diabetic Therapeutic Strategies. Cells. 2024;13(15). Epub 20240724. doi: 10.3390/cells13151244. PubMed PMID: 39120275; PMCID: PMC11311556.

17. Eng J, Kleinman WA, Singh L, Singh G, Raufman JP. Isolation and characterization of exendin-4, an exendin-3 analogue, from Heloderma suspectum venom. Further evidence for an exendin receptor on dispersed acini from guinea pig pancreas. J Biol Chem. 1992;267(11):7402–5. PubMed PMID: 1313797.

18. Olofsson CS, Gopel SO, Barg S, Galvanovskis J, Ma X, Salehi A, Rorsman P, Eliasson L. Fast insulin secretion reflects exocytosis of docked granules in mouse pancreatic B-cells. Pflugers Arch. 2002;444(1-2):43–51. Epub 20020131. doi: 10.1007/s00424-002-0781-5. PubMed PMID: 11976915.

19. Binder JX, Pletscher-Frankild S, Tsafou K, Stolte C, O’Donoghue SI, Schneider R, Jensen LJ. COMPARTMENTS: unification and visualization of protein subcellular localization evidence. Database (Oxford). 2014;2014:bau012. Epub 20140225. doi: 10.1093/database/bau012. PubMed PMID: 24573882; PMCID: PMC3935310.

20. Phang HQ, Hoon JL, Lai SK, Zeng Y, Chiam KH, Li HY, Koh CG. POPX2 phosphatase regulates the KIF3 kinesin motor complex. J Cell Sci. 2014;127(Pt 4):727–39. Epub 20131211. doi: 10.1242/jcs.126482. PubMed PMID: 24338362.

21. Xiong R, Rao P, Kim S, Li M, Wen X, Yuan W. Herpes Simplex Virus 1 US3 Phosphorylates Cellular KIF3A To Downregulate CD1d Expression. J Virol. 2015;89(13):6646–55. Epub 20150415. doi: 10.1128/JVI.00214-15. PubMed PMID: 25878107; PMCID: PMC4468489.

22. Ichinose S, Ogawa T, Hirokawa N. Mechanism of Activity-Dependent Cargo Loading via the Phosphorylation of KIF3A by PKA and CaMKIIa. Neuron. 2015;87(5):1022–35. doi: 10.1016/j.neuron.2015.08.008. PubMed PMID: 26335646.

23. Goodwin SS, Vale RD. Patronin regulates the microtubule network by protecting microtubule minus ends. Cell. 2010;143(2):263–74. doi: 10.1016/j.cell.2010.09.022. PubMed PMID: 20946984; PMCID: PMC3008421.

24. Wu Y, Foollee A, Chan AY, Hille S, Hauke J, Challis MP, Johnson JL, Yaron TM, Mynard V, Aung OH, Cleofe MAS, Huang C, Lim Kam Sian TCC, Rahbari M, Gallage S, Heikenwalder M, Cantley LC, Schittenhelm RB, Formosa LE, Smith GC, Okun JG, Muller OJ, Rusu PM, Rose AJ. Phosphoproteomics-directed manipulation reveals SEC22B as a hepatocellular signaling node governing metabolic actions of glucagon. Nat Commun. 2024;15(1):8390. Epub 20240927. doi: 10.1038/s41467-024-52703-w. PubMed PMID: 39333498; PMCID: PMC11436942.

25. Dalle S. Targeting Protein Kinases to Protect Beta-Cell Function and Survival in Diabetes. Int J Mol Sci. 2024;25(12). Epub 20240611. doi: 10.3390/ijms25126425. PubMed PMID: 38928130; PMCID: PMC11203834.

26. Jiang WJ, Peng YC, Yang KM. Cellular signaling pathways regulating beta-cell proliferation as a promising therapeutic target in the treatment of diabetes. Exp Ther Med. 2018;16(4):3275–85. Epub 20180813. doi: 10.3892/etm.2018.6603. PubMed PMID: 30233674; PMCID: PMC6143874.

27. Hornbeck PV, Kornhauser JM, Tkachev S, Zhang B, Skrzypek E, Murray B, Latham V, Sullivan M. PhosphoSitePlus: a comprehensive resource for investigating the structure and function of experimentally determined post-translational modifications in man and mouse. Nucleic Acids Res. 2012;40(Database issue):D261–70. Epub 20111201. doi: 10.1093/nar/gkr1122. PubMed PMID: 22135298; PMCID: PMC3245126.

28. Ercu M, Klussmann E. Roles of A-Kinase Anchoring Proteins and Phosphodiesterases in the Cardiovascular System. J Cardiovasc Dev Dis. 2018;5(1). Epub 20180220. doi: 10.3390/jcdd5010014. PubMed PMID: 29461511; PMCID: PMC5872362.

29. Calejo AI, Tasken K. Targeting protein-protein interactions in complexes organized by A kinase anchoring proteins. Front Pharmacol. 2015;6:192. Epub 20150908. doi: 10.3389/fphar.2015.00192. PubMed PMID: 26441649; PMCID: PMC4562273.

30. Falcone JI, Scott JD. The ascent of AKAPs, from architectural elements to kinase anchors: a perspective. Biochem J. 2025;482(10):485–98. Epub 20250513. doi: 10.1042/BCJ20253085. PubMed PMID: 40364611; PMCID: PMC12203939.

31. Schillace RV, Scott JD. Organization of kinases, phosphatases, and receptor signaling complexes. J Clin Invest. 1999;103(6):761–5. doi: 10.1172/JCI6491. PubMed PMID: 10079095; PMCID: PMC408155.

32. Ahmad F, Murata T, Shimizu K, Degerman E, Maurice D, Manganiello V. Cyclic nucleotide phosphodiesterases: important signaling modulators and therapeutic targets. Oral Dis. 2015;21(1):e25–50. Epub 20140912. doi: 10.1111/odi.12275. PubMed PMID: 25056711; PMCID: PMC4275405.

33. Degerman E, Ahmad F, Chung YW, Guirguis E, Omar B, Stenson L, Manganiello V. From PDE3B to the regulation of energy homeostasis. Curr Opin Pharmacol. 2011;11(6):676–82. Epub 20111014. doi: 10.1016/j.coph.2011.09.015. PubMed PMID: 22001403; PMCID: PMC3225700.

34. Cygnar KD, Zhao H. Phosphodiesterase 1C is dispensable for rapid response termination of olfactory sensory neurons. Nat Neurosci. 2009;12(4):454–62. Epub 20090322. doi: 10.1038/nn.2289. PubMed PMID: 19305400; PMCID: PMC2712288.

35. Debruyne DN, Bhatnagar N, Sharma B, Luther W, Moore NF, Cheung NK, Gray NS, George RE. ALK inhibitor resistance in ALK(F1174L)-driven neuroblastoma is associated with AXL activation and induction of EMT. Oncogene. 2016;35(28):3681–91. Epub 20151130. doi: 10.1038/onc.2015.434. PubMed PMID: 26616860; PMCID: PMC4885798.

36. Zhang T, Inesta-Vaquera F, Niepel M, Zhang J, Ficarro SB, Machleidt T, Xie T, Marto JA, Kim N, Sim T, Laughlin JD, Park H, LoGrasso PV, Patricelli M, Nomanbhoy TK, Sorger PK, Alessi DR, Gray NS. Discovery of potent and selective covalent inhibitors of JNK. Chem Biol. 2012;19(1):140–54. doi: 10.1016/j.chembiol.2011.11.010. PubMed PMID: 22284361; PMCID: PMC3270411.

37. Zhou W, Ercan D, Chen L, Yun CH, Li D, Capelletti M, Cortot AB, Chirieac L, Iacob RE, Padera R, Engen JR, Wong KK, Eck MJ, Gray NS, Janne PA. Novel mutant-selective EGFR kinase inhibitors against EGFR T790M. Nature. 2009;462(7276):1070–4. doi: 10.1038/nature08622. PubMed PMID: 20033049; PMCID: PMC2879581.

38. Drewes G, Ebneth A, Preuss U, Mandelkow EM, Mandelkow E. MARK, a novel family of protein kinases that phosphorylate microtubule-associated proteins and trigger microtubule disruption. Cell. 1997;89(2):297–308. doi: 10.1016/s0092-8674(00)80208-1. PubMed PMID: 9108484.

39. Timm T, Li XY, Biernat J, Jiao J, Mandelkow E, Vandekerckhove J, Mandelkow EM. MARKK, a Ste20-like kinase, activates the polarity-inducing kinase MARK/PAR-1. EMBO J. 2003;22(19):5090–101. doi: 10.1093/emboj/cdg447. PubMed PMID: 14517247; PMCID: PMC204455.

40. Fye MA, Kaverina I. Insulin secretion hot spots in pancreatic beta cells as secreting adhesions. Front Cell Dev Biol. 2023;11:1211482. Epub 20230526. doi: 10.3389/fcell.2023.1211482. PubMed PMID: 37305687; PMCID: PMC10250740.

41. Yu SB, Pekkurnaz G. Mechanisms Orchestrating Mitochondrial Dynamics for Energy Homeostasis. J Mol Biol. 2018;430(21):3922–41. Epub 20180805. doi: 10.1016/j.jmb.2018.07.027. PubMed PMID: 30089235; PMCID: PMC6186503.

42. Chen W, Zhao H, Li Y. Mitochondrial dynamics in health and disease: mechanisms and potential targets. Signal Transduct Target Ther. 2023;8(1):333. Epub 20230906. doi: 10.1038/s41392-023-01547-9. PubMed PMID: 37669960; PMCID: PMC10480456.

43. Scott I, Youle RJ. Mitochondrial fission and fusion. Essays Biochem. 2010;47:85–98. doi: 10.1042/bse0470085. PubMed PMID: 20533902; PMCID: PMC4762097.

44. Severin M, Pedersen EL, Borre MT, Axholm I, Christiansen FB, Ponniah M, Czaplinska D, Larsen T, Pardo LA, Pedersen SF. Dynamic localization of the Na+-HCO3-co-transporter NBCn1 to the plasma membrane, centrosomes, spindle and primary cilia. J Cell Sci. 2023;136(7). Epub 20230411. doi: 10.1242/jcs.260687. PubMed PMID: 37039101.

45. Zhu X, Hu R, Brissova M, Stein RW, Powers AC, Gu G, Kaverina I. Microtubules Negatively Regulate Insulin Secretion in Pancreatic beta Cells. Dev Cell. 2015;34(6):656–68. doi: 10.1016/j.devcel.2015.08.020. PubMed PMID: 26418295; PMCID: PMC4594944.

46. Shida T, Cueva JG, Xu Z, Goodman MB, Nachury MV. The major alpha-tubulin K40 acetyltransferase alphaTAT1 promotes rapid ciliogenesis and efficient mechanosensation. Proc Natl Acad Sci U S A. 2010;107(50):21517–22. Epub 20101110. doi: 10.1073/pnas.1013728107. PubMed PMID: 21068373; PMCID: PMC3003046.

47. Rao Y, Hao R, Wang B, Yao TP. A Mec17-Myosin II Effector Axis Coordinates Microtubule Acetylation and Actin Dynamics to Control Primary Cilium Biogenesis. PLoS One. 2014;9(12):e114087. Epub 20141210. doi: 10.1371/journal.pone.0114087. PubMed PMID: 25494100; PMCID: PMC4262394.

48. Kalebic N, Sorrentino S, Perlas E, Bolasco G, Martinez C, Heppenstall PA. alphaTAT1 is the major alpha-tubulin acetyltransferase in mice. Nat Commun. 2013;4:1962. doi: 10.1038/ncomms2962. PubMed PMID: 23748901.

49. Trogden KP, Lee J, Bracey KM, Ho KH, McKinney H, Zhu X, Arpag G, Folland TG, Osipovich AB, Magnuson MA, Zanic M, Gu G, Holmes WR, Kaverina I. Microtubules regulate pancreatic beta-cell heterogeneity via spatiotemporal control of insulin secretion hot spots. Elife. 2021;10. Epub 20211116. doi: 10.7554/eLife.59912. PubMed PMID: 34783306; PMCID: PMC8635970.

50. Trogden KP, Zhu X, Lee JS, Wright CVE, Gu G, Kaverina I. Regulation of Glucose-Dependent Golgi-Derived Microtubules by cAMP/EPAC2 Promotes Secretory Vesicle Biogenesis in Pancreatic beta Cells. Curr Biol. 2019;29(14):2339–50 e5. Epub 20190711. doi: 10.1016/j.cub.2019.06.032. PubMed PMID: 31303487; PMCID: PMC6698911.

51. Li L, Yang XJ. Tubulin acetylation: responsible enzymes, biological functions and human diseases. Cell Mol Life Sci. 2015;72(22):4237–55. Epub 20150731. doi: 10.1007/s00018-015-2000-5. PubMed PMID: 26227334; PMCID: PMC11113413.

52. Deb Roy A, Gross EG, Pillai GS, Seetharaman S, Etienne-Manneville S, Inoue T. Non-catalytic allostery in alpha-TAT1 by a phospho-switch drives dynamic microtubule acetylation. J Cell Biol. 2022;221(11). Epub 20221012. doi: 10.1083/jcb.202202100. PubMed PMID: 36222836; PMCID: PMC9565784.

53. Iuzzolino A, Pellegrini FR, Rotili D, Degrassi F, Trisciuoglio D. The alpha-tubulin acetyltransferase ATAT1: structure, cellular functions, and its emerging role in human diseases. Cell Mol Life Sci. 2024;81(1):193. Epub 20240423. doi: 10.1007/s00018-024-05227-x. PubMed PMID: 38652325; PMCID: PMC11039541.

54. Hormozdiari F, van de Bunt M, Segre AV, Li X, Joo JWJ, Bilow M, Sul JH, Sankararaman S, Pasaniuc B, Eskin E. Colocalization of GWAS and eQTL Signals Detects Target Genes. Am J Hum Genet. 2016;99(6):1245–60. Epub 20161117. doi: 10.1016/j.ajhg.2016.10.003. PubMed PMID: 27866706; PMCID: PMC5142122.

55. Nicolae DL, Gamazon E, Zhang W, Duan S, Dolan ME, Cox NJ. Trait-associated SNPs are more likely to be eQTLs: annotation to enhance discovery from GWAS. PLoS Genet. 2010;6(4):e1000888. Epub 20100401. doi: 10.1371/journal.pgen.1000888. PubMed PMID: 20369019; PMCID: PMC2848547.

56. Turn RE, Aziz-Zanjani MO, Asthana A, Jackson PK. Strategies for multimodal spatiotemporal profiling of phosphorylation in cilia biology. J Cell Sci. 2025;138(20). Epub 20251031. doi: 10.1242/jcs.264159. PubMed PMID: 41171145.

57. Harbour JW, Dean DC. The Rb/E2F pathway: expanding roles and emerging paradigms. Genes Dev. 2000;14(19):2393–409. doi: 10.1101/gad.813200. PubMed PMID: 11018009.

58. Mittnacht S, Lees JA, Desai D, Harlow E, Morgan DO, Weinberg RA. Distinct sub-populations of the retinoblastoma protein show a distinct pattern of phosphorylation. EMBO J. 1994;13(1):118–27. doi: 10.1002/j.1460-2075.1994.tb06241.x. PubMed PMID: 8306955; PMCID: PMC394785.

59. Spiering MJ. The discovery of GABA in the brain. J Biol Chem. 2018;293(49):19159–60. doi: 10.1074/jbc.CL118.006591. PubMed PMID: 30530855; PMCID: PMC6295731.

60. Metherate R, Ashe JH. GABAergic suppression prevents the appearance and subsequent fatigue of an NMDA receptor-mediated potential in neocortex. Brain Res. 1995;699(2):221–30. doi: 10.1016/0006-8993(95)00909-a. PubMed PMID: 8616625.

61. Lai S, Pelech S. Regulatory roles of conserved phosphorylation sites in the activation T-loop of the MAP kinase ERK1. Mol Biol Cell. 2016;27(6):1040–50. Epub 20160128. doi: 10.1091/mbc.E15-07-0527. PubMed PMID: 26823016; PMCID: PMC4791125.

62. Wang Z, Thurmond DC. Mechanisms of biphasic insulin-granule exocytosis - roles of the cytoskeleton, small GTPases and SNARE proteins. J Cell Sci. 2009;122(Pt 7):893–903. doi: 10.1242/jcs.034355. PubMed PMID: 19295123; PMCID: PMC2720925.

63. Chen S, Owens GC, Makarenkova H, Edelman DB. HDAC6 regulates mitochondrial transport in hippocampal neurons. PLoS One. 2010;5(5):e10848. Epub 20100526. doi: 10.1371/journal.pone.0010848. PubMed PMID: 20520769; PMCID: PMC2877100.

64. Even A, Morelli G, Broix L, Scaramuzzino C, Turchetto S, Gladwyn-Ng I, Le Bail R, Shilian M, Freeman S, Magiera MM, Jijumon AS, Krusy N, Malgrange B, Brone B, Dietrich P, Dragatsis I, Janke C, Saudou F, Weil M, Nguyen L. ATAT1-enriched vesicles promote microtubule acetylation via axonal transport. Sci Adv. 2019;5(12):eaax2705. Epub 20191218. doi: 10.1126/sciadv.aax2705. PubMed PMID: 31897425; PMCID: PMC6920029.

65. David Y, Castro IG, Schuldiner M. The Fast and the Furious: Golgi Contact Sites. Contact (Thousand Oaks). 2021;4:1–15. Epub 20210809. doi: 10.1177/25152564211034424. PubMed PMID: 35071979; PMCID: PMC7612241.

66. Chen B, Lyssiotis CA, Shah YM. Mitochondria-organelle crosstalk in establishing compartmentalized metabolic homeostasis. Mol Cell. 2025;85(8):1487–508. doi: 10.1016/j.molcel.2025.03.003. PubMed PMID: 40250411; PMCID: PMC12182711.

67. Hanada K. Lipid transfer proteins rectify inter-organelle flux and accurately deliver lipids at membrane contact sites. J Lipid Res. 2018;59(8):1341–66. Epub 20180608. doi: 10.1194/jlr.R085324. PubMed PMID: 29884707; PMCID: PMC6071762.

68. Nagashima S, Tabara LC, Tilokani L, Paupe V, Anand H, Pogson JH, Zunino R, McBride HM, Prudent J. Golgi-derived PI(4)P-containing vesicles drive late steps of mitochondrial division. Science. 2020;367(6484):1366–71. doi: 10.1126/science.aax6089. PubMed PMID: 32193326.

69. Doyle CP, Timple L, Hammond GRV. OSBP is a Major Determinant of Golgi Phosphatidylinositol 4-Phosphate Homeostasis. Contact (Thousand Oaks). 2024;7:25152564241232196. Epub 20240222. doi: 10.1177/25152564241232196. PubMed PMID: 38405037; PMCID: PMC10893830.

70. Camunas-Soler J, Dai XQ, Hang Y, Bautista A, Lyon J, Suzuki K, Kim SK, Quake SR, MacDonald PE. Patch-Seq Links Single-Cell Transcriptomes to Human Islet Dysfunction in Diabetes. Cell Metab. 2020;31(5):1017–31 e4. Epub 20200416. doi: 10.1016/j.cmet.2020.04.005. PubMed PMID: 32302527; PMCID: PMC7398125.

71. Parkes DG, Pittner R, Jodka C, Smith P, Young A. Insulinotropic actions of exendin-4 and glucagon-like peptide-1 in vivo and in vitro. Metabolism. 2001;50(5):583–9. doi: 10.1053/meta.2001.22519. PubMed PMID: 11319721.

72. Kolterman OG, Kim DD, Shen L, Ruggles JA, Nielsen LL, Fineman MS, Baron AD. Pharmacokinetics, pharmacodynamics, and safety of exenatide in patients with type 2 diabetes mellitus. Am J Health Syst Pharm. 2005;62(2):173–81. doi: 10.1093/ajhp/62.2.173. PubMed PMID: 15700891.

73. Maselli DB, Camilleri M. Effects of GLP-1 and Its Analogs on Gastric Physiology in Diabetes Mellitus and Obesity. Adv Exp Med Biol. 2021;1307:171–92. doi: 10.1007/5584_2020_496. PubMed PMID: 32077010.

74. Johnson JL, Yaron TM, Huntsman EM, Kerelsky A, Song J, Regev A, Lin TY, Liberatore K, Cizin DM, Cohen BM, Vasan N, Ma Y, Krismer K, Robles JT, van de Kooij B, van Vlimmeren AE, Andree-Busch N, Kaufer NF, Dorovkov MV, …, Cantley LC. An atlas of substrate specificities for the human serine/threonine kinome. Nature. 2023;613(7945):759–66. Epub 20230111. doi: 10.1038/s41586-022-05575-3. PubMed PMID: 36631611; PMCID: PMC9876800.

