## Supplementary figures and images for "Metabolic STAMP reveals GPCR signaling networks that program β-cell insulin secretion"

### Fig.S1

#### Overview of metabolic STAMP

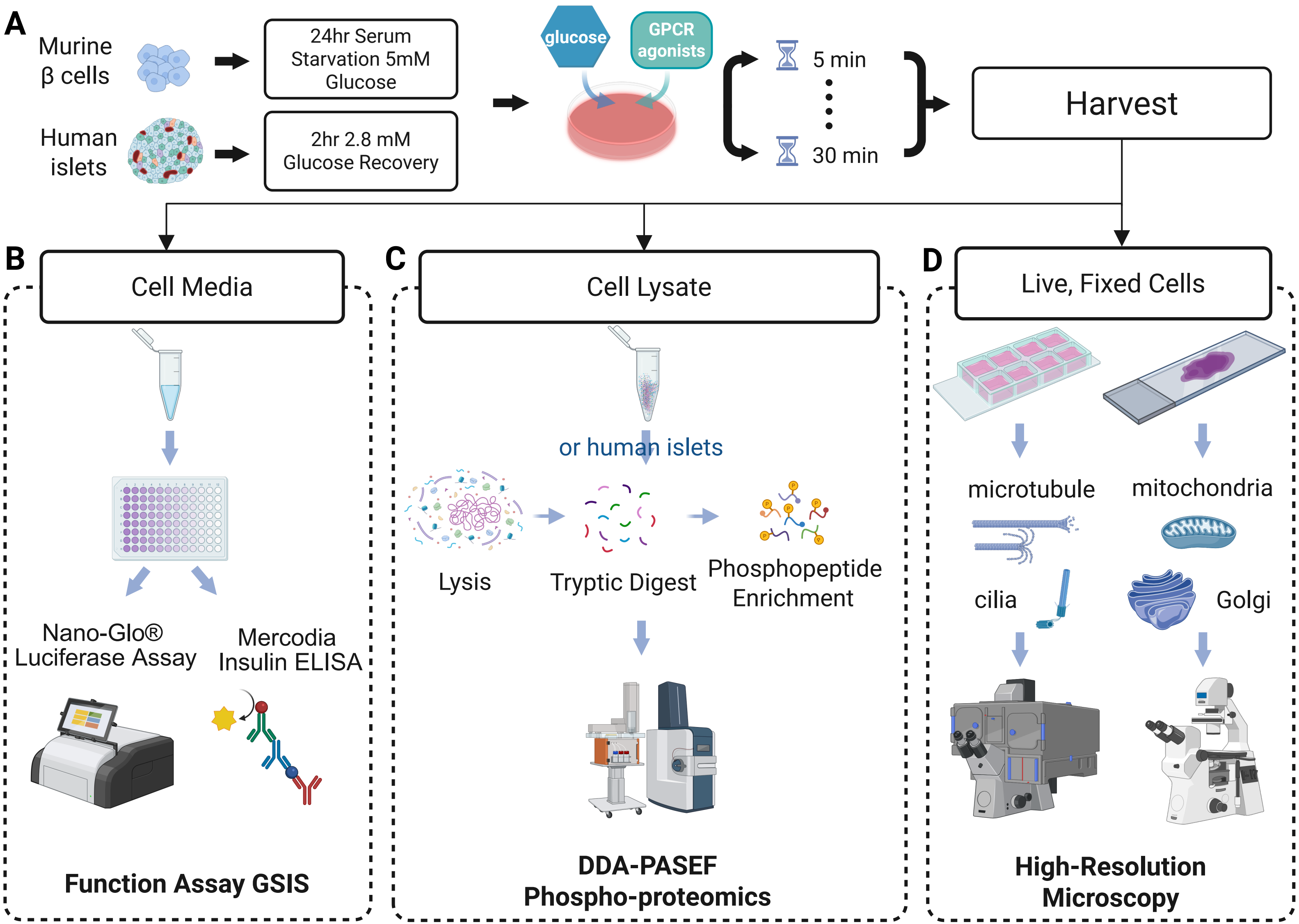

### Fig.S2

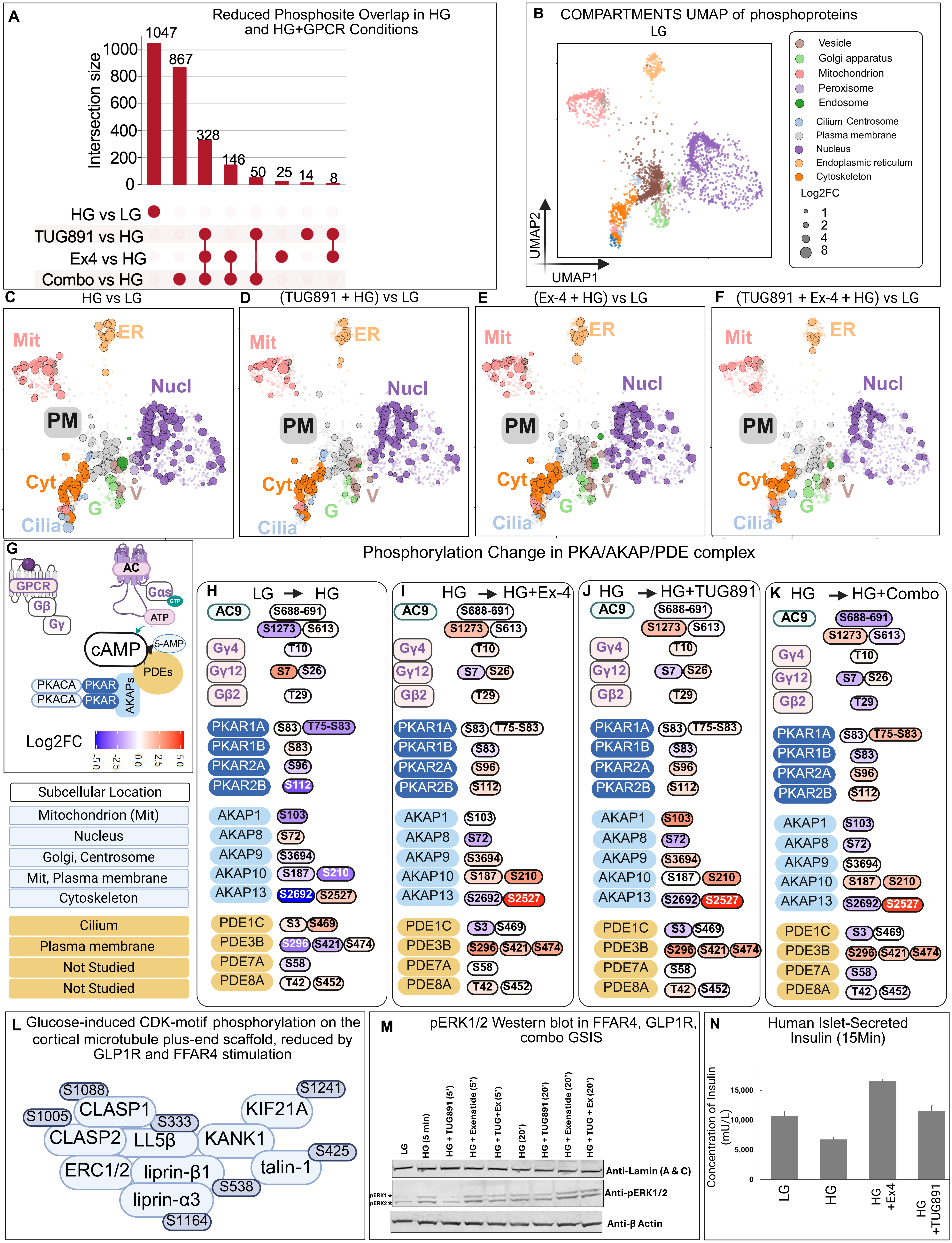

### Fig.S4

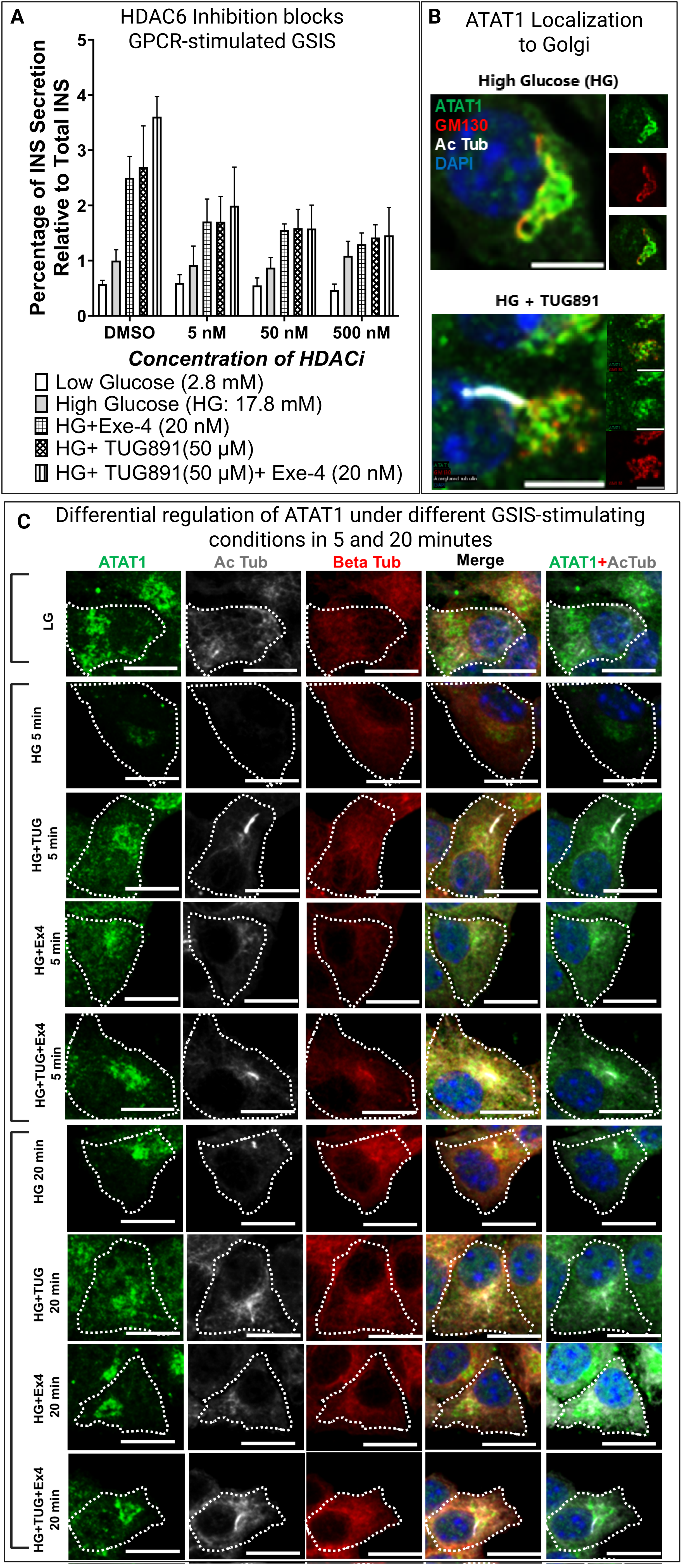

### Fig.S6

#### Distribution of Acetylated Tubulin in Pancreatic Islet α Cells

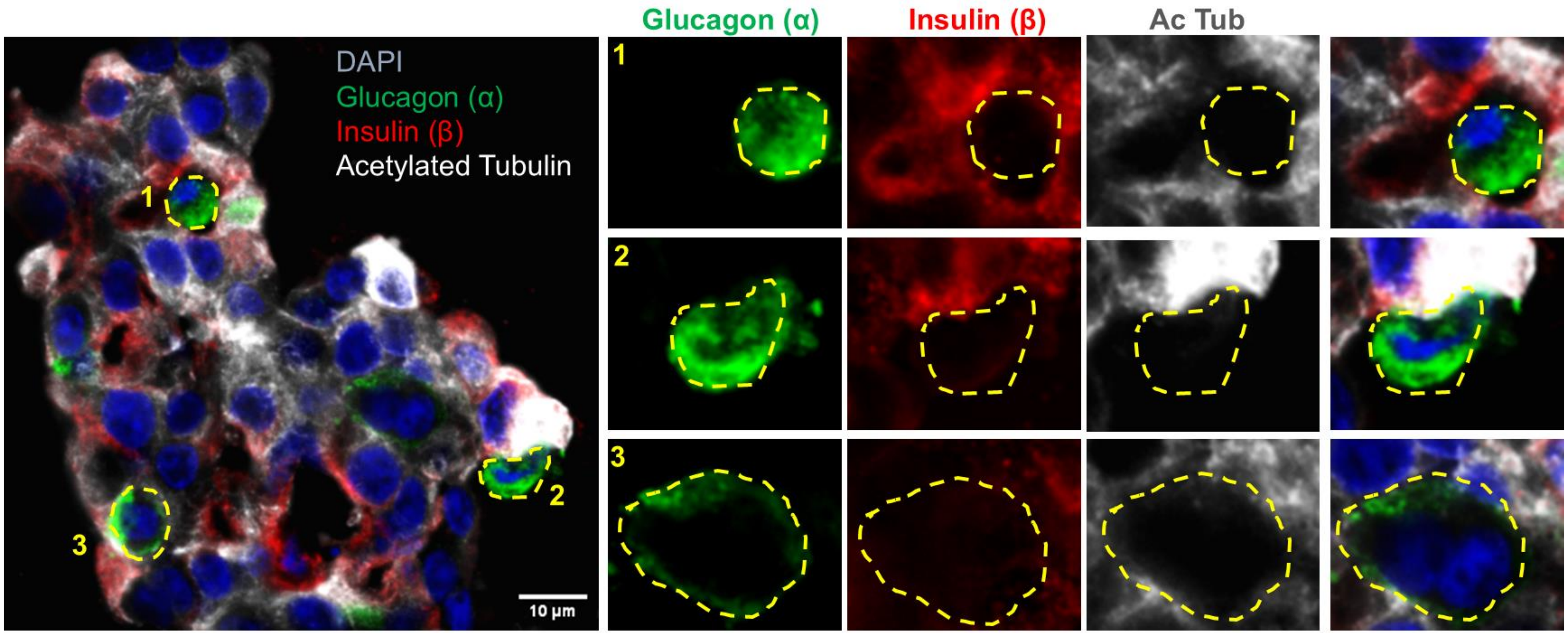
