## Supplementary material for "Metabolic STAMP reveals GPCR signaling networks that program β-cell insulin secretion": Fig.S3

### Workflow for kinase inhibitor screening with phosphoproteomics to identify kinase–substrate relationships

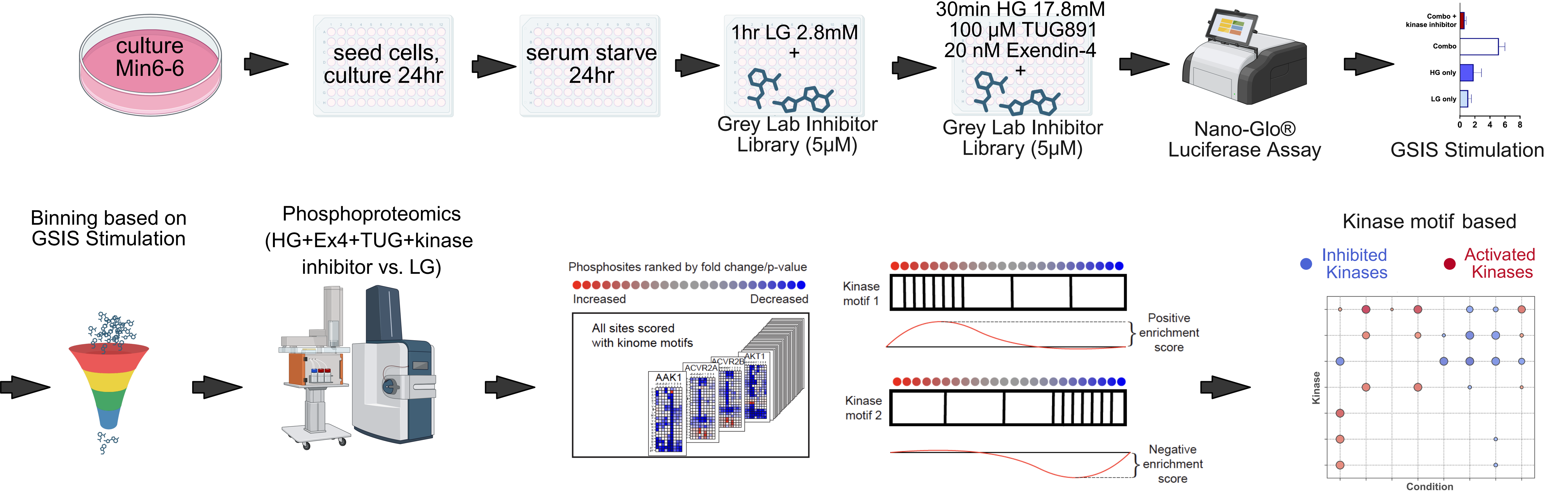
