## Supplementary material for "Metabolic STAMP reveals GPCR signaling networks that program β-cell insulin secretion": Fig.S5

### ATAT1 nuclear distribution shows differential localization based on GSIS stimulatory conditions

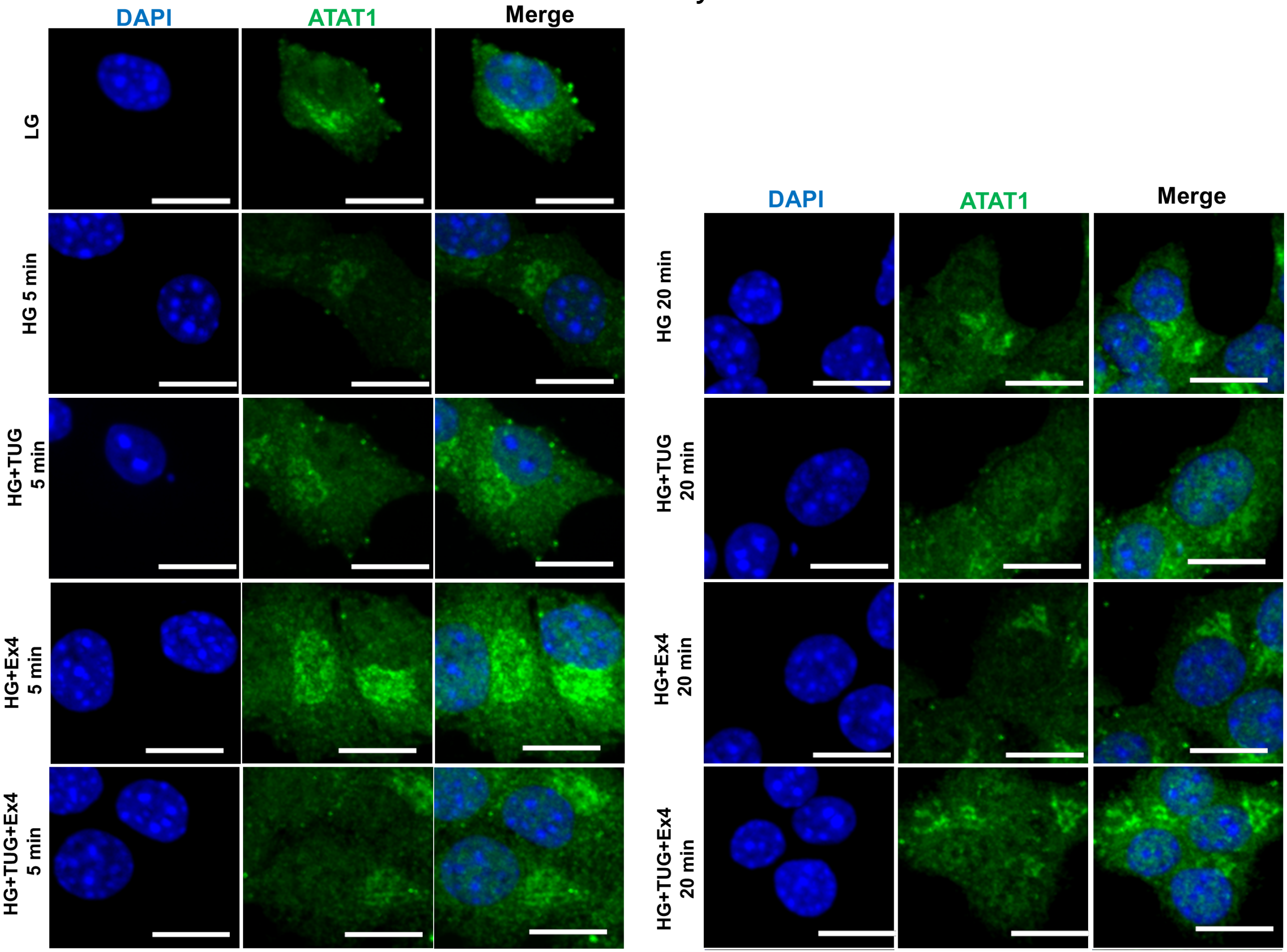
