## Supplementary material for "Metabolic STAMP reveals GPCR signaling networks that program β-cell insulin secretion": Table S4

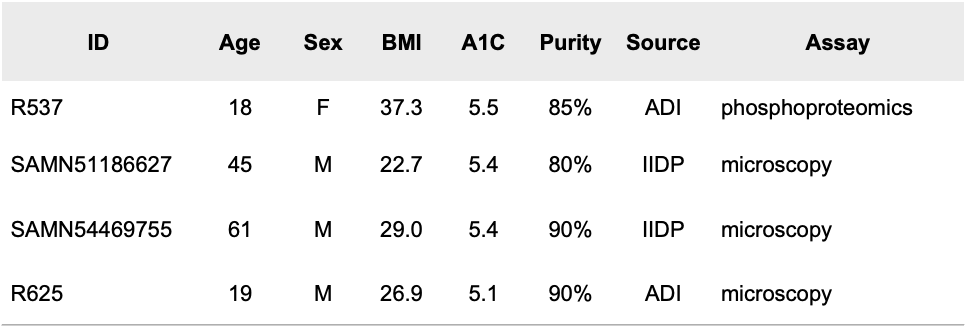

IIDP, Integrated Islet Distribution Program.

ADI, Alberta Diabetes Institute.

**Table S5. Human Islets Information**
